# Cooperative Regulation of Membrane Domain Registration by Lipid Headgroup Properties and Cholesterol Dynamics

**DOI:** 10.64898/2026.01.16.699701

**Authors:** Jinyu Chen, Yu Cao, D. Peter Tieleman, Qing Liang

**Author notes:** These authors contributed equally (Jinyu Chen, Yu Cao).

## Abstract

The lateral organization of lipid domains and their coupling across the two leaflets are central to the structure of plasma membranes, yet the molecular determinants of domain registration and anti-registration remain incompletely understood. Here, we employ coarse-grained molecular dynamics simulations to investigate how lipid headgroup properties and cholesterol (CHOL) concentration regulate domain registration and anti-registration in symmetric model bilayers. Using a free energy perturbation (FEP)-like headgroup-interpolation approach to continuously interpolate between headgroup representations characteristic of phosphatidylethanolamine (PE) and phosphatidylcholine (PC) lipids at a fixed tail composition, we show that headgroup properties and CHOL concentration act cooperatively through their effects on lipid packing and membrane curvature. Simulations with restrained CHOL flip-flop further demonstrate that transbilayer CHOL exchange actively strengthens interleaflet coupling and promotes domain registration. These results identify headgroup-dependent packing, CHOL-induced curvature, and CHOL flip-flop as cooperative determinants of domain organization in model membranes.

**TOC Graphic:** 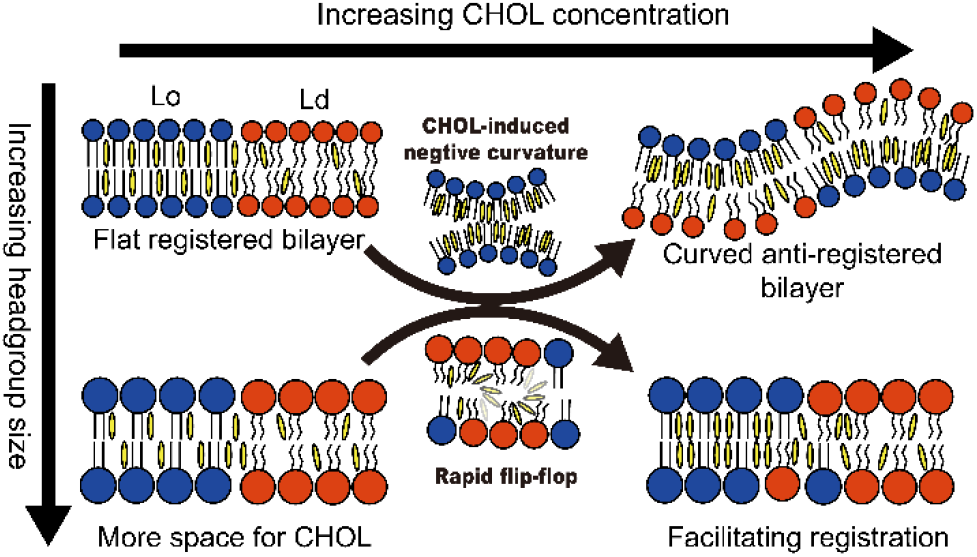

## Introduction

Plasma membranes of cells are highly heterogeneous and asymmetric systems composed of diverse proteins and lipids.^1–3^ The lipid raft hypothesis proposes that plasma membranes contain dynamic lateral heterogeneities enriched in cholesterol and sphingolipids, which have been implicated in cellular processes such as signal transduction and selective molecular transport.^4,5^ However, the lateral and transbilayer organization of plasma membranes remains incompletely understood because of their structural and compositional complexity.^6–8^ Model membranes with simplified compositions and architectures have therefore been widely used to investigate the physical principles underlying membrane organization.^9,10^

In phase-separated model membranes, lipid domains in opposing leaflets adopt either registered configurations, in which like phases are aligned across the bilayer, or anti-registered configurations, in which unlike phases oppose one another.^11–13^ Considerable effort has been devoted to elucidating the molecular determinants underlying these structures. Experimental studies have demonstrated the existence of interleaflet coupling and have identified lipid intrinsic curvature and composition as key factors modulating the coupling strength.^14–17^ Complementary simulation and theoretical investigations have revealed that domain registration and anti-registration are influenced by multiple factors, including hydrophobic mismatch, the positions of double bonds in lipid tails, membrane bending rigidity and undulations, and domain boundary line tension.^11,13,18–22^

Specifically, cholesterol (CHOL), a principal sterol component of mammalian plasma membranes, serves as a critical regulator of membrane structure and function.^3,23–26^ Most recently, Doktorova et al. demonstrated that CHOL sustains interleaflet phospholipid asymmetry through rapid transbilayer flip-flop, accumulating in the phospholipid-depleted exoplasmic leaflet where it preferentially interacts with saturated sphingolipids while buffering differential leaflet stress.^3^ Furthermore, CHOL flip-flop enhances interleaflet coupling between coexisting lipid domains of the same phase and promotes domain registration, with midplane CHOL molecules (i.e., those residing at the bilayer midplane) contributing significantly to this phenomenon.^27–29^

Many proposed mechanisms of interleaflet coupling involve interactions within the hydrophobic region of the bilayer, and previous studies have therefore focused primarily on the effects of lipid tails on domain registration and anti-registration.^11,19^ In comparison, the potential influence of lipid headgroups on the interleaflet coupling of the lipid domains has received much less attention. Particularly, how CHOL and lipid headgroup properties jointly regulate lipid domain registration and anti-registration remains elusive.

In this work, we employ coarse-grained molecular dynamics simulations together with a free energy perturbation (FEP)-like headgroup-interpolation approach to systematically examine how lipid headgroup properties and CHOL concentration regulate domain registration and anti-registration. By continuously varying the lipid headgroup properties while maintaining a fixed lipid tail composition, we isolate the contribution of the headgroups from that of the hydrophobic tails. We find that increasing CHOL concentration favors anti-registration more strongly in bilayers composed of lipids with smaller and more strongly interacting headgroups, whereas bilayers composed of lipids with larger and more weakly interacting headgroups remain relatively flat and registered over a broader range of CHOL concentrations. This difference arises from distinct responses of lipid packing and membrane curvature to CHOL incorporation. Additional simulations in which CHOL flip-flop is selectively restrained show that transbilayer CHOL exchange actively contributes to interleaflet coupling and domain registration. These results reveal a cooperative mechanism involving headgroup-dependent packing, CHOL-induced curvature, and CHOL flip-flop.

## Model and Methods

### System Setup

As summarized in Table 1, two types of lipid bilayers with distinct compositions were constructed to investigate the effects of CHOL concentration and lipid headgroup properties on the registration and anti-registration of lipid domains. One system consists of dipalmitoyl-phosphatidylcholine (DPPC), dilinoleoyl-PC (DIPC), and CHOL, whereas the other comprises dipalmitoyl-phosphatidylethanolamine (DPPE), dilinoleoyl-PE (DIPE), and CHOL. For each system, the CHOL concentration was varied among 12.5, 30, and 50 mol %, while maintaining a constant molar ratio between the other two lipid components. These lipid compositions were chosen as model systems to systematically isolate the effect of headgroup chemistry (PC vs. PE) at fixed tail composition; they are not intended to reproduce the compositional complexity of biological plasma membranes.

**Table 1:** Overview of Simulation Systems.

| System | Lipid molar ratio | # lipids per leaflet <sup>a</sup> | Duration | # water molecules |
| --- | --- | --- | --- | --- |
| **DPPC/DIPC/CHOL** |  |  |  |  |
| I | 4:3:1 | 760:570:190 | $2 \times 10 \mu\text{s}$ | 74,851 |
| II | 4:3:3 | 608:456:456 | $2 \times 10 \mu\text{s}$ | 75,873 |
| III | 4:3:7 | 434:325:760 | $2 \times 10 \mu\text{s}$ | 77,039 |
| **DPPE/DIPE/CHOL** |  |  |  |  |
| IV | 4:3:1 | 760:570:190 | $2 \times 10 \mu\text{s}$ | 74,851 |
| V | 4:3:3 | 608:456:456 | $2 \times 10 \mu\text{s}$ | 75,873 |
| VI | 4:3:7 | 434:325:760 | $2 \times 10 \mu\text{s}$ | 77,079 |

<sup>a</sup> Initial structures of all lipid bilayers are symmetric, with the identical number of each lipid type in both leaflets.

As shown in Figure 1a, DPPC and DPPE are fully saturated lipids, whereas DIPC and DIPE are unsaturated, each containing two cis double bonds in their acyl chains. PC and PE lipids differ primarily in their headgroup properties, with PC having a larger effective headgroup and PE being capable of hydrogen bonding, owing to differences in their chemical structures. Figure 1b illustrates the initial structures of two lipid bilayers, composed of DPPC/DIPC/CHOL and DPPE/DIPE/CHOL, respectively. Previous studies have shown that such ternary lipid bilayers can spontaneously phase separate into liquid-ordered (Lo) and liquid-disordered (Ld) domains under comparable conditions,^5,13–15,29,30^ supporting their use as model systems for investigating lipid domain registration and anti-registration.

**Figure 1.**
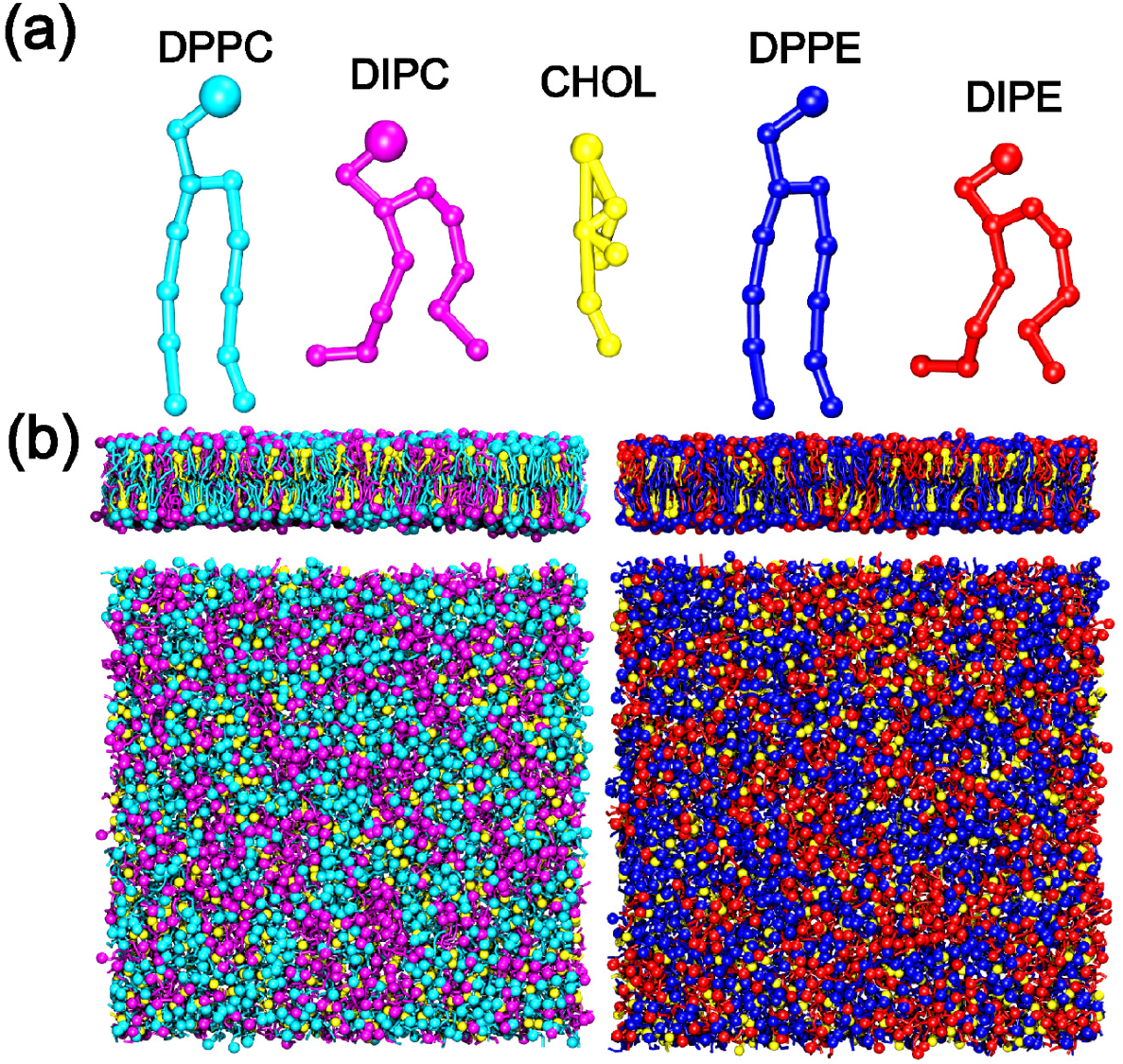
(a) Coarse-grained structures of DPPC (cyan), DIPC (magenta), CHOL (yellow), DPPE (blue), and DIPE (red). This color scheme is used consistently throughout the paper. (b) Lateral and top views of the initial lipid bilayers composed of DPPC, DIPC, and CHOL (left) and of DPPE, DIPE, and CHOL (right), shown for Systems II and V as examples.

It should be noted that the present study employs symmetric ternary lipid bilayers described by a coarse-grained representation. Although such models have been widely adopted to elucidate the fundamental principles governing lipid domain organization,^10,11^ they do not capture the compositional asymmetry, lipid heterogeneity, and the presence of membrane proteins of the plasma membrane. Thus, the results reported here should be interpreted as mechanistic insights derived from simplified model membranes, which nonetheless provide a useful framework for understanding lipid domain organization in more complex cellular membranes.

All lipid bilayers were built using the script INSANE,^31^ and the compositions of the lipid bilayers are shown in Table 1. The simulation box size was set large enough to minimize the potential influence of finite size on the phase separation of the lipid bilayers, with the initial size of 30 × 30 × 15 nm^3^. Additionally, 0.15 M sodium chloride (NaCl) was added to mimic the physiological conditions. Particularly, in coarse-grained simulations, the parameters of *LINCS order* of 12 and *LINCS iter* of 2 were used for CHOL molecules to avoid artificial temperature gradients due to non-convergent constraints.^32^

### Simulation Details

All MD simulations were performed using the GROMACS 2021.4 package, employing the coarse-grained MARTINI v2.2 force field to represent lipids, water molecules, and ions.^33–35^ Van der Waals (vdW) interactions were treated with a cutoff distance of 1.2 nm, with the Lennard-Jones potential smoothly shifted to zero between 0.9 and 1.2 nm. Electrostatic interactions were described using the Coulombic potential with a cutoff of 1.2 nm and the default relative dielectric constant of 15. All simulations were conducted in the isothermal-isobaric (*NpT*) ensemble. A pressure of 1 bar was applied separately in the bilayer plane (*xy*-plane) and along the normal (*z*) direction using a semi-isotropic Parrinello-Rahman pressure coupling scheme, ^36^ with a coupling constant of 5 ps and a compressibility of 3 × 10⁻⁴ bar⁻¹. Temperature was maintained at 310 K using the velocity-rescaling coupling scheme^37^ with a time constant of 1 ps, with separate thermostats for the lipid bilayer and solvent. Periodic boundary conditions were applied in all three directions. Each simulation was run for 10 μs with a time step of 20 fs, following energy minimization and equilibration.

### FEP-like Simulations for Examining the Effect of Lipid Headgroup Properties

In the coarse-grained MARTINI model, the headgroups of PC and PE are represented by Q0 and Qd beads, respectively. The two bead types differ in both van der Waals radius (effective size) and hydrogen-bonding capability, implemented through distinct Lennard-Jones interaction parameters with other bead types (Qd acts as a hydrogen-bond donor, while Q0 does not). To systematically study the impact of lipid headgroup properties on the lipid domain registration and anti-registration, we gradually varied the vdW radius of the lipid headgroup from Q0 to Qd using a free energy perturbation (FEP)-like method.^38–41^ Lipids with Q0 and Qd headgroups were treated as two distinct states of the same lipid species. The transition from the Qd state (PE-like) to the Q0 state (PC-like) was achieved by tuning a coupling parameter λ from 0 to 1, which modulated vdW interactions analogously to the standard FEP method,^42^

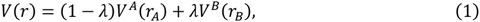

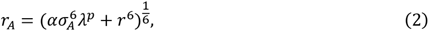

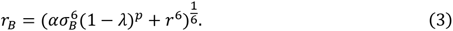

Here, *V^A^* and *V^B^* represent the standard van der Waals (vdW) potentials corresponding to the Qd state (*λ* = 0) and the Q0 state (*λ* = 1), respectively. The parameter *α* denotes the coupling strength of the potential, and *σ* represents the interaction radius. Since Q0 and Qd differ in both van der Waals radius and hydrogen-bonding capability, the coupling parameter λ captures the combined effect of effective headgroup size and headgroup-mediated interactions. To examine the effect of lipid headgroup properties, eight replica systems were constructed with *λ* values of 0, 0.2, 0.4, 0.5, 0.6, 0.7, 0.8, and 1. The parameters *α* and *p* were set to 0.5 and 1, respectively, for all replicas.

### Simulations with Cholesterol Flip-Flop Restraints

To investigate the effect of CHOL flip-flop on membrane domain registration and anti-registration, we performed several simulations with the flip-flop of CHOL restrained by introducing a short-ranged repulsive potential between the ROH bead of CHOL and the terminal bead of tail chains of lipids.^43^ These results were then compared to those from normal simulations where CHOL flip-flop was unrestrained.

### Data Analysis

MD simulation trajectories were mainly analyzed using GROMACS 2021.4, Visual Molecular Dynamics (VMD),^44^ LiPyphilic,^45^ MDAnalysis,^46^ and MemSurfer.^47^ MDAnalysis was used for trajectory processing and molecular selections. The domain registration fraction (DRF) and cholesterol (CHOL) flip-flop were calculated using LiPyphilic, and the area per lipid (APL) was calculated using MemSurfer.

The DRF was defined as the Pearson correlation coefficient between the two-dimensional (2D) number-density distributions of ordered-domain lipid markers in the upper and lower leaflets:

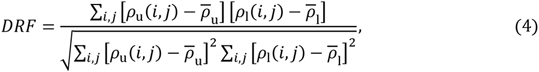

where *ρ*_u_ and *ρ*_l_ are the upper- and lower-leaflet densities, respectively. The DRF values of 1 and −1 indicate complete registration and complete anti-registration, respectively. Leaflet assignment was performed using the *AssignCurvedLeaflets* module in LiPyphilic,^45^ which identifies the two leaflets from the three-dimensional connectivity of lipid beads rather than from a fixed z-coordinate. The GL1 and GL2 beads of non-sterol lipids were grouped using a connectivity cutoff of 12 Å under periodic boundary conditions. CHOL was assigned independently using its ROH bead with a cutoff of 10 Å. The 2D density distributions were constructed from the PO4 beads of saturated phospholipids and the ROH beads of CHOL. The default LiPyphilic grid resolution of approximately 1 Å was used, and the density functions were smoothed with a periodic Gaussian kernel with a standard deviation of 12 Å before calculating the Pearson correlation coefficient.

APL was calculated separately for the two leaflets using MemSurfer,^47^ which reconstructed separate periodic three-dimensional triangulated surfaces for the upper and lower leaflets using one representative headgroup bead per lipid. The APL of lipid *i* was calculated as the mixed-Voronoi area associated with its surface vertex:

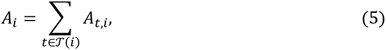

where T(*i*) denotes the surface triangles containing vertex *i*, and *A_t,i_* is the portion of triangle *t* assigned to that vertex. Thus, the calculated APL accounts for the three-dimensional curvature of each leaflet, and the APL values were averaged over both leaflets.

Lo-domain surface areas and CHOL number densities were calculated separately for the upper and lower leaflets over 8–10 μs. Lo regions were identified on the MemSurfer-reconstructed leaflet surfaces from the local enrichment of saturated relative to unsaturated lipid headgroups within 12 Å, and regions with positive saturated-lipid enrichment were classified as Lo domains. All Lo domains in each leaflet were included. The Lo-domain surface area was summed over these regions, and the CHOL number density was calculated as the number of leaflet-assigned CHOL molecules within them divided by their total surface area.

Lipid-tail order parameters were calculated over 5–10 μs. For tail bond *b* of lipid *i* at time *t*,

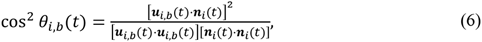

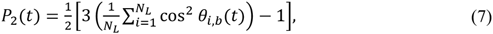

where ***u****_i,b_*(*t*) is the vector between the two beads defining tail bond *b*, ***n****_i_*(*t*) is the corresponding local membrane normal calculated using MemSurfer,^47^ and *N_L_* is the number of lipids of that type. For each of the eight reported bonds, the order parameter was averaged over 5–10 μs, after excluding the first three headgroup/linker bonds.

Midplane CHOL molecules were defined as CHOL molecules whose ROH beads were not within 12 Å of the representative headgroup beads of either leaflet. Their numbers were averaged over 5–10 μs and divided by the total number of CHOL molecules to obtain the midplane CHOL fraction.

CHOL flip-flop was analyzed using the *FlipFlop* module in LiPyphilic.^45^ Each CHOL molecule was classified as belonging to the upper leaflet, lower leaflet, or bilayer midplane according to the position of its ROH bead. A transition was counted as successful only when the CHOL molecule reached the opposite leaflet and remained there for at least 100 ns, a criterion that excludes transient excursions across the midplane. The flip-flop rate per CHOL molecule was calculated as:

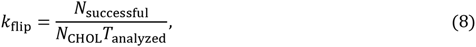

where *N*_successful_ is the number of successful transitions, *N*_CHOL_ is the total number of CHOL molecules, and *T*_analyzed_ is the duration of the analyzed trajectory segment (the final 5 μs of each simulation).

## Results and Discussion

### Lipid Domain Registration and Anti-registration in PC and PE bilayers

We first investigated the phase separation of six ternary lipid bilayers composed of PC (or PE) and CHOL at different molar ratios (Table 1). Snapshots of the phase separation processes are shown in Figures S1–S6 in the Supporting Information (SI), and the final structures of the bilayers are presented in Figure 2. In PC bilayers (Figure 2a–c), lipid domains exhibit registered structures in the lipid bilayers with 12.5 mol% and 30 mol% CHOL, whereas domain anti-registration occurs in the lipid bilayers with 50 mol% CHOL. However, PE bilayers (Figure 2d–f) display domain registration only at the lower cholesterol concentration of 12.5 mol%, with anti-registration observed at the higher concentrations of 30 and 50 mol%. Notably, lipid domain anti-registration is typically associated with increased membrane curvature.

**Figure 2.**
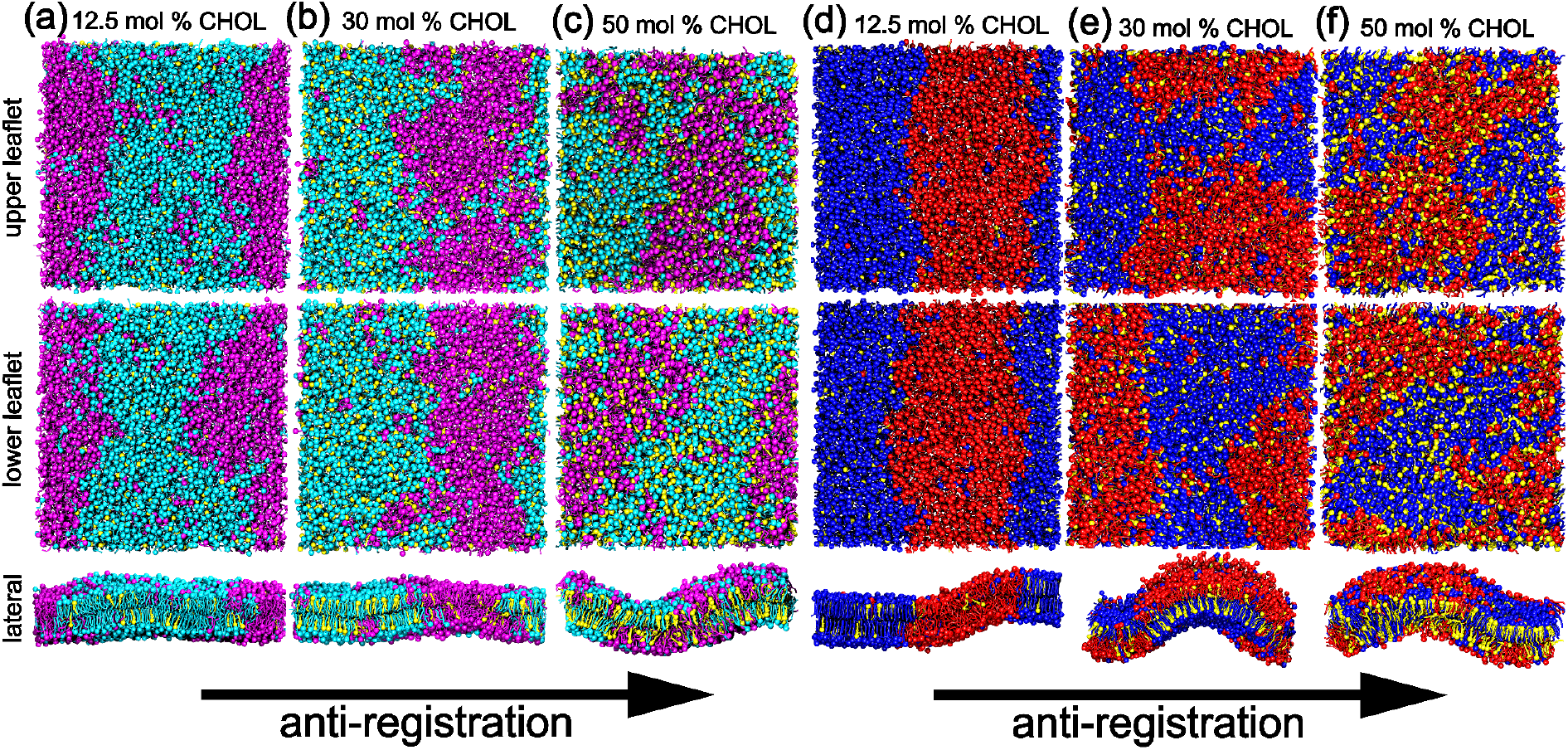
Snapshots of final structures of lipid bilayers at the end of 10 µs simulations for: (a) System I: PC bilayer with 12.5 mol% CHOL, (b) System II: PC bilayer with 30 mol% CHOL, (c) System III: PC bilayer with 50 mol% CHOL, (d) System IV: PE bilayer with 12.5 mol% CHOL, (e) System V: PE bilayer with 30 mol% CHOL, and (f) System VI: PE bilayer with 50 mol% CHOL. For each system, the top, middle, and bottom rows correspond to the upper leaflet, lower leaflet, and lateral view, respectively.

Although lipids with different headgroups respond differently to changes in CHOL concentration, both PC and PE lipid bilayers gradually transition from registration to anti-registration with increasing CHOL concentration. Since PC and PE lipids in our coarse-grained representation differ primarily in headgroup properties (effective size and hydrogen-bonding capability) while sharing identical tail structures, these results indicate that lipid headgroup properties and CHOL concentration jointly influence the registration and anti-registration of lipid domains.

### Cooperative Effects of CHOL Concentration and Headgroup Properties on Domain Registration and Anti-registration

To systematically elucidate the effects of CHOL concentration and lipid headgroup properties on domain registration and anti-registration, we employed a FEP-like approach to gradually transform PE-like headgroups into PC-like headgroups by adjusting the coupling parameter λ. Conceptually, a larger λ corresponds to a more PC-like headgroup with a larger effective size and weaker interactions. This was performed in lipid bilayers containing CHOL at 12.5 mol% and 30 mol% (see Model and Methods for details). The domain registration fraction (DRF), defined in Model and Methods, was used to characterize the degree of registration and anti-registration, as shown in Figure 3.

**Figure 3.**
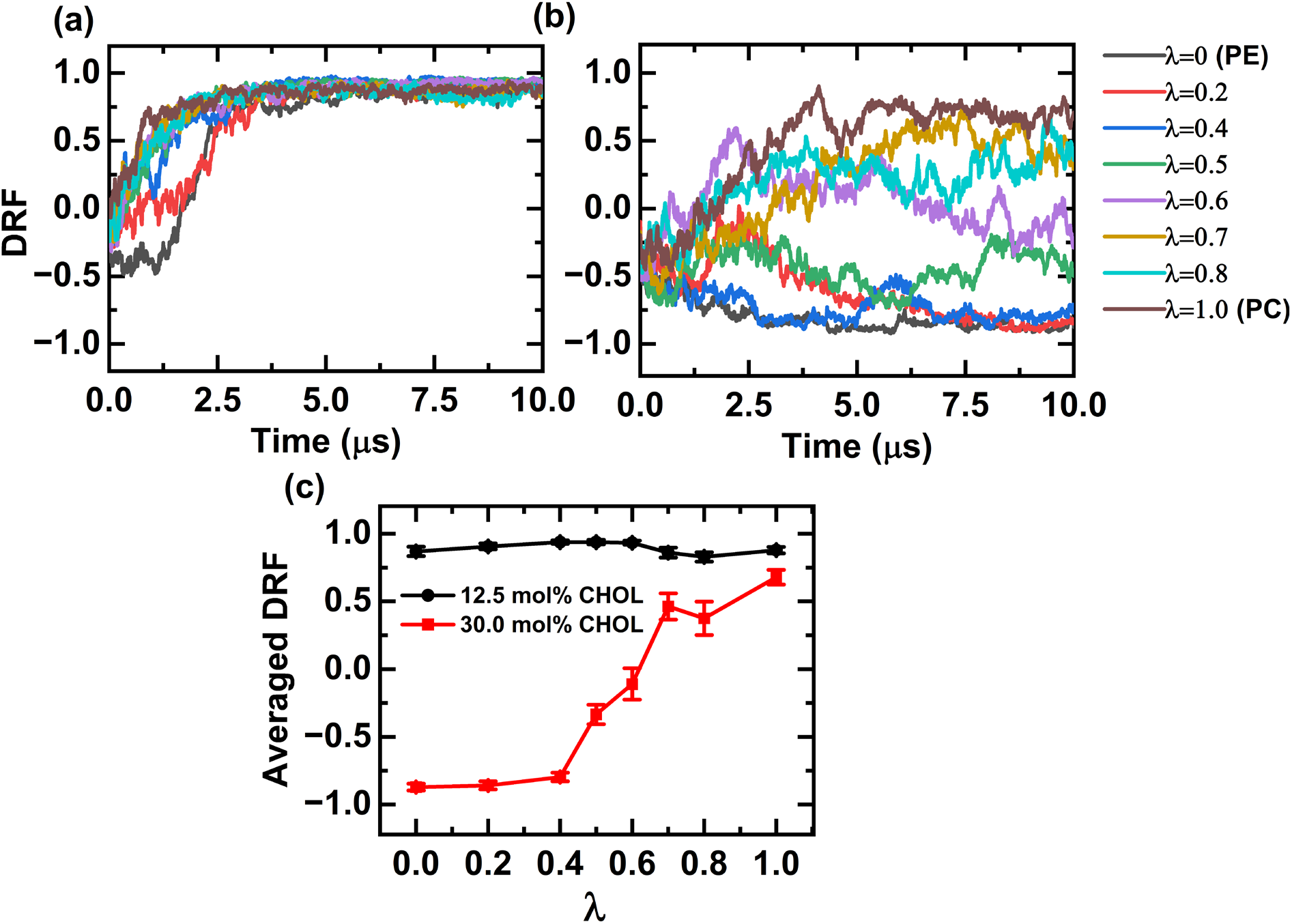
(a, b) Domain registration fraction (DRF) of lipid bilayers with CHOL concentrations of 12.5 mol% and 30 mol%, respectively, shown as a function of simulation time for different λ values representing varying lipid headgroup properties. (c) Time-averaged DRFs of lipid bilayers at 12.5 mol% and 30 mol% CHOL, calculated over the last 2 μs of each simulation, as a function of λ.

For the lipid bilayers with 12.5 mol% CHOL, the DRFs of all eight systems increase from smaller values (approximately −0.5 to 0) to larger values (∼0.75) within the initial 5 µs and stabilize between 0.8 and 0.9 during the subsequent 5 µs, as shown in Figure 3a. This indicates that the transbilayer domain organization reaches a stable registered state within approximately 5 µs and that lipid domains are registered in all eight systems. Therefore, at this lower CHOL concentration (12.5 mol%), the effect of lipid headgroup properties is not explicitly manifested in domain registration versus anti-registration.

As the CHOL concentration increases to 30 mol%, the DRFs vary dramatically as the parameter λ changes between 0 and 1, as shown in Figure 3b,c. At relatively small λ values (0, 0.2, and 0.4), corresponding to PE-like lipids with smaller and strongly interacting headgroups, the bilayer DRFs remain predominantly below −0.5 throughout the 10 µs simulations, indicating that lipid domains in these bilayers are anti-registered for most of the simulation time. At intermediate λ values of 0.5 and 0.6, the bilayer DRFs fluctuate within the range of −0.5 to 0.25, with average values of approximately −0.25 and 0 over the last 2 µs (Figure 3c), respectively, making it difficult to clearly classify the domains as registered or anti-registered. However, with increasing λ, the DRFs increase substantially, suggesting a trend toward domain registration. At relatively large λ values (0.7–1.0), corresponding to PC-like lipids with larger and weakly interacting headgroups, the bilayer DRFs exceed 0.75, indicating that lipid domains are registered. These results demonstrate that lipid headgroup properties play a significant role in determining domain registration and anti-registration only when the CHOL concentration is sufficiently high, as shown in Figure 3c.

Furthermore, given that CHOL molecules preferentially localize within the Lo domains enriched in saturated lipids,^5,14,15,29,48^ we calculated the APL of saturated lipids in bilayers containing 12.5 mol% and 30 mol% CHOL using FEP-like simulations with varying λ over the simulation time as shown in Figure 4. In both systems, the APL of saturated lipids increases with increasing λ, indicating that lipid headgroup properties play an important role in determining lipid packing within the Lo domain.

**Figure 4.**
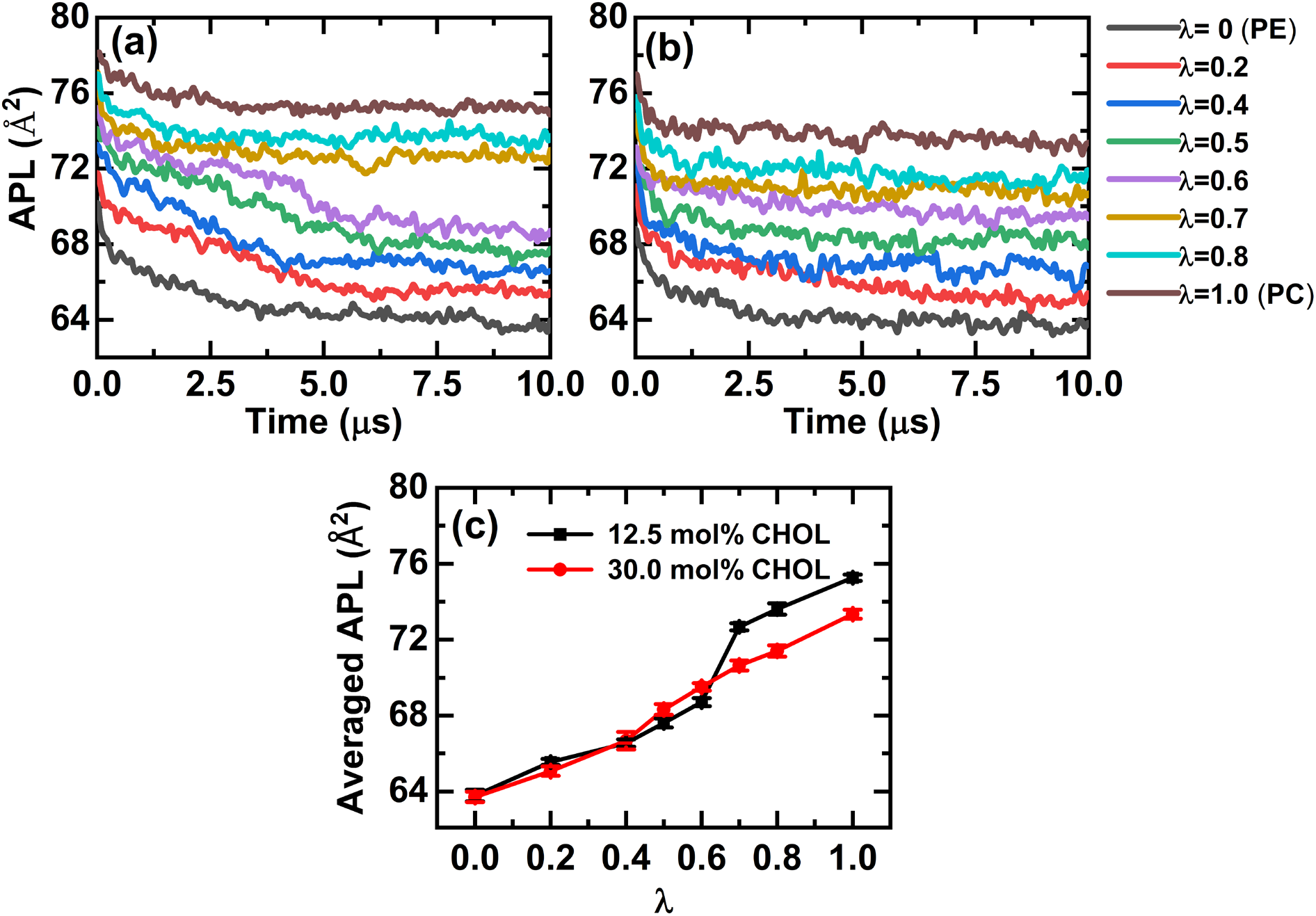
(a, b) Area per lipid (APL) of saturated lipids in Lo domains of lipid bilayers containing 12.5 mol% and 30 mol% CHOL, respectively, shown as a function of simulation time for different λ values. (c) Time-averaged APL of saturated lipids at the two CHOL concentrations, calculated over the last 2 μs of each simulation, as a function of λ.

Notably, the APL of PE-like bilayers (*λ* ≤ 0.4) is nearly identical at the two CHOL concentrations (Figure 4c), despite the much higher CHOL number density within the Lo domains at 30 mol% CHOL (Figure S7). In PC-like bilayers (λ ≥ 0.7), by contrast, the APL at 30 mol% is measurably lower than at 12.5 mol% (Figure 4c), consistent with the condensing effect of CHOL.^25–27^ This difference indicates that the two headgroup types accommodate additional CHOL through different structural pathways.

In the Lo domain, the highly ordered packing of lipid tails constrains the lateral spacing between neighboring lipids, which is in turn modulated by headgroup size and headgroup– headgroup interactions. The packing perturbation introduced by CHOL may therefore be accommodated either through changes in the lateral packing and interfacial area of the headgroups, or through changes in lipid-tail orientation, order, and local packing. PE headgroups are small and form strong, compact interactions that constrain interfacial rearrangement. In these bilayers, the packing perturbation is therefore accommodated primarily through changes in tail orientation and order. PC headgroups are larger and interact more weakly, allowing the interfacial area to adjust directly and thereby respond through a more pronounced change in APL while largely preserving tail order.

In agreement with this, tail order parameters computed relative to local membrane normals change substantially for DPPE between 12.5 and 30 mol% CHOL, whereas the corresponding DPPC values change only weakly (Figure S8). Because local normals were used, this response is not a trivial consequence of global membrane bending. Similarly, since the APL was computed on the reconstructed three-dimensional membrane surface, the nearly constant APL in PE-like systems is not a projection artifact. The unchanged APL and the pronounced tail-order response in PE-like systems are two complementary manifestations of the same underlying packing behavior, and do not indicate that PE-like Lo domains are insensitive to increased CHOL content. Both headgroup types respond to added CHOL, but the response is partitioned differently between the headgroup interface and the hydrophobic core. This picture is consistent with experimental observations that PE headgroups exhibit stronger intermolecular association,^49^ that progressive methylation of PE toward PC alters headgroup size and hydrogen-bonding capacity,^50^ and that PE bilayers have smaller molecular areas and more ordered acyl chains than PC bilayers with identical tails.^51^

On this basis, we hypothesize that a CHOL saturation concentration exists within the Lo domain, operationally defined as the concentration above which the packing perturbation can no longer be accommodated through interfacial rearrangement and bilayer curvature develops. This is a phenomenological definition based on our simulation observations rather than a rigorous thermodynamic quantity. This threshold is higher for PC-like Lo domains, where the larger and more weakly associated headgroups provide both greater interfacial space and greater capacity for rearrangement, and lower for PE-like Lo domains. This threshold therefore sets the CHOL concentration at which a given bilayer transitions from registered to anti-registered domains.

At 12.5 mol% CHOL, the concentration is below this threshold in both PC and PE bilayers, and the domains remain registered (Figures 2 and 3). At 30 mol%, PC bilayers still accommodate the added CHOL through interfacial rearrangement and maintain a flat, registered morphology. In PE-like and intermediate bilayers (λ < 0.7), however, 30 mol% likely exceeds the threshold: the strongly interacting headgroups resist lateral expansion while the effective volume of the CHOL-enriched hydrophobic region increases, producing a volumetric imbalance between the tail and headgroup layers. This imbalance induces a tendency toward negative curvature in the Lo domains of both leaflets. To preserve bilayer integrity, once a registered Lo domain in one leaflet becomes negatively curved, the corresponding Lo domain in the opposing leaflet is replaced by a Ld domain with positive curvature, converting a flat bilayer with registered domains into a curved bilayer with anti-registered domains (Figures 2 and 3).

With the CHOL concentration increasing to 50 mol%, both PC and PE bilayers exhibit anti-registered domains (Figures 2c and f), providing additional support for this mechanism. Previous theoretical work has proposed that bilayer bending modulates the elastic energy of the Lo domain and its boundary, effectively lowering the line tension that stabilizes registered configurations.^21^ Although we do not directly calculate line tension here, our observation that anti-registration coincides with the development of bilayer curvature is qualitatively consistent with this framework. Thus, headgroup properties, together with the CHOL saturation concentration, cooperatively determine how the CHOL-induced packing perturbation is partitioned between the headgroup interface and the hydrophobic core, thereby regulating Lo domain curvature and promoting anti-registration.

### Role of CHOL Flip-Flop in Lipid Domain Registration and Anti-registration

We further investigated the role of transbilayer CHOL flip-flop in regulating lipid domain registration and anti-registration. The time-averaged CHOL flip-flop rates, calculated over the last 5 µs of the FEP-like simulations for lipid bilayers containing 12.5 mol% and 30 mol% CHOL, are shown in Figure 5. At 12.5 mol% CHOL, the flip-flop rate increases modestly from (3.17±0.28) ×10^3^ s^−1^ to (4.38±0.25) ×10^3^ s^−1^ with increasing parameter λ. At 30 mol% CHOL, however, the flip-flop rate increases much more dramatically from (6.01±0.44) ×10^2^ s^−1^ to (2.54±0.07) ×10^3^ s^−1^ as λ increases. Furthermore, as shown in Figure 4c, the APL of saturated lipids also increases with increasing λ. These findings suggest that CHOL flip-flop dynamics are strongly influenced by the packing environment of lipid tails, particularly within the Lo domain.

**Figure 5.**
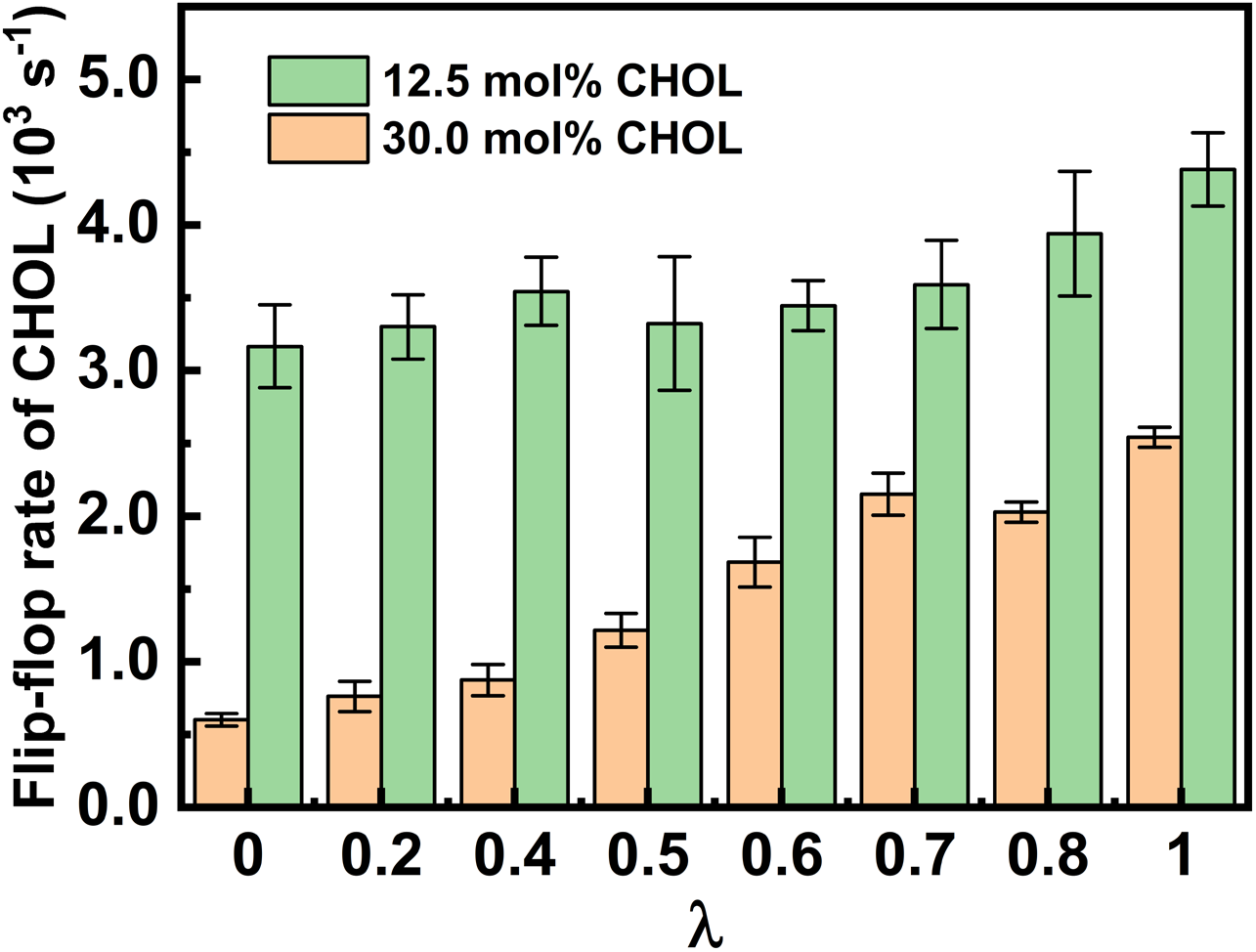
Time-averaged flip-flop rate per CHOL molecule as a function of λ in eight FEP-like simulation replicas at two CHOL concentrations (12.5 mol% and 30 mol%). Error bars indicate the standard deviation of the measured flip-flop rates over the last 5 µs of each simulation.

Lipids with a larger APL exhibit larger transient inter-lipid spacing and density fluctuations, which facilitate the flip-flop of CHOL molecules. In the bilayer with 12.5 mol% CHOL, the concentration remains below the saturation threshold, so transient density fluctuations readily accommodate CHOL translocation. Consequently, the CHOL flip-flop rate is relatively high and increases slightly with increasing λ, as shown in Figure 5.

At 30 mol% CHOL, by contrast, the concentration likely exceeds the saturation threshold, particularly in systems composed of lipids with smaller headgroups, thereby reducing the transient density fluctuations that permit CHOL translocation. As a result, CHOL flip-flop is strongly suppressed in Lo domains enriched in saturated lipids with smaller headgroups (λ≤0.6). As λ increases, the inter-lipid spacing increases correspondingly, providing greater transient volume for CHOL translocation and leading to a notable increase in the flip-flop rate (Figure 5).

Furthermore, comparing Figures 3 and 5 reveals a positive correlation between the CHOL flip-flop rate and domain registration. In principle, flip-flop could induce an asymmetric transbilayer distribution of CHOL and thereby influence domain organization. However, our simulations show no measurable imbalance in the number of CHOL molecules between the two leaflets; the populations in opposing leaflets remain equal over 8–10 μs (Figure S7). Flip-flop therefore appears to affect domain registration in a dynamic rather than a static manner.

In bilayers with 12.5 mol% CHOL, the relatively rapid flip-flop coincides with registered Lo domains in both PC-like and PE-like bilayers. At 30 mol% CHOL, the behavior becomes strongly dependent on headgroup properties: PE-like bilayers (λ ≤ 0.6) show substantially reduced flip-flop rates (Figure 5) together with anti-registered domains, whereas PC-like bilayers (λ > 0.6) retain faster flip-flop and registered configurations (Figure 3). This correlation suggests that flip-flop contributes to interleaflet coupling within Lo domains, a possibility we test directly below using simulations with restrained flip-flop.

To test whether CHOL flip-flop actively contributes to interleaflet coupling, we performed additional simulations on the four reference systems containing 12.5 and 30 mol% CHOL (Systems I, II, IV, and V in Table 1), in which CHOL flip-flop was restrained by introducing a short-ranged repulsive interaction between ROH beads of CHOL molecules and the terminal beads of lipid tails. The time-averaged CHOL flip-flop rates, calculated over the last 5 µs of these simulations, indicate that the restraint reduces the rate by approximately one to two orders of magnitude (Figure 6).

**Figure 6.**
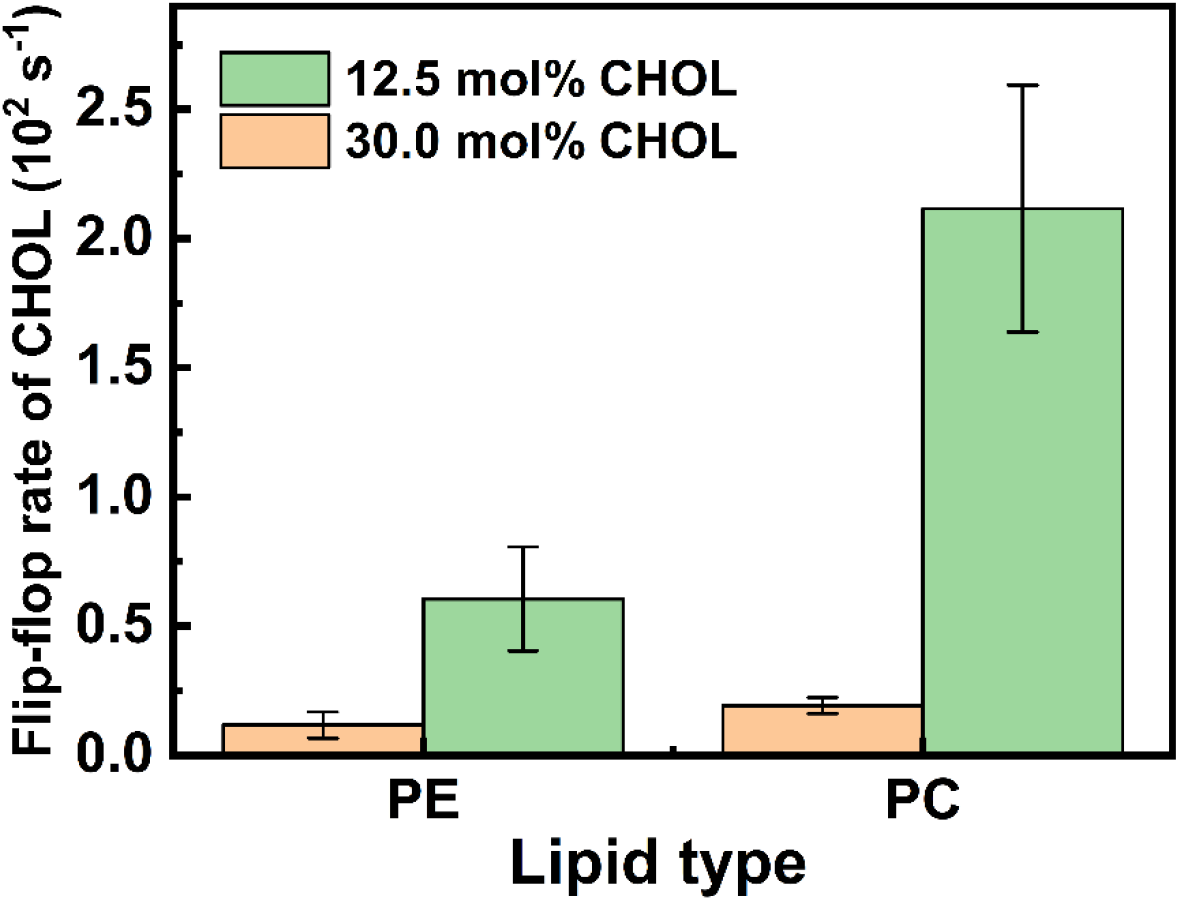
Time-averaged flip-flop rate per CHOL molecule in four lipid bilayers composed of PC and PE at CHOL concentrations of 12.5 mol% and 30 mol%, with CHOL flip-flop restrained. Error bars indicate standard deviations calculated over the last 5 µs of each simulation.

The DRFs of the CHOL-restrained systems are shown in Figure 7, together with those of the corresponding CHOL-unrestrained systems for comparison. For bilayers with 12.5 mol% CHOL (Figure 7a), restraining flip-flop leads to a slight decrease in the DRF of the PC bilayer, while the lipid domains remain registered as shown in Figure S9. For the PE bilayer at the same CHOL concentration, the DRF decreases dramatically and fluctuates around zero. The corresponding simulation snapshots (Figure S10) indicate the coexistence of registered and anti-registered domains in the CHOL-restrained PE bilayer.

**Figure 7.**
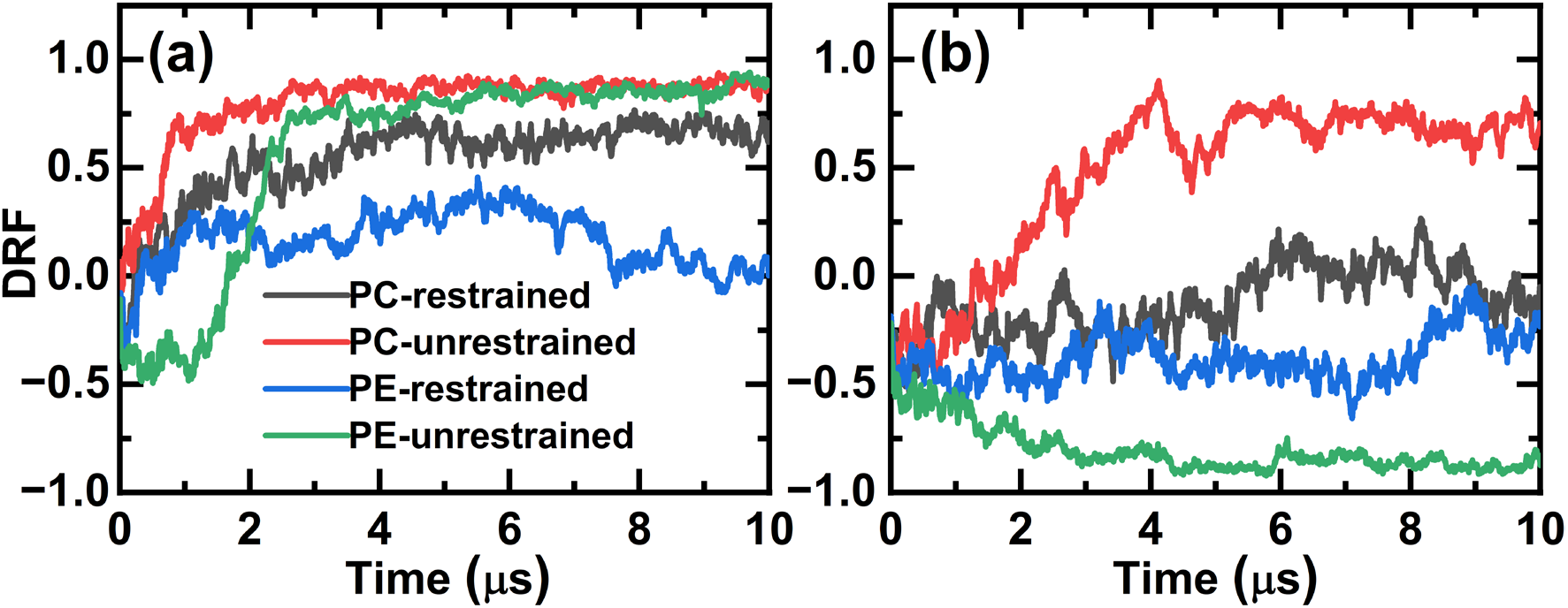
Domain registration fraction (DRF) as a function of simulation time for lipid bilayers with restrained and unrestrained CHOL flip-flop at CHOL concentrations of (a) 12.5 mol% and (b) 30 mol%. Data are shown for PC bilayers with restrained (black) and unrestrained (red) flip-flop, and for PE bilayers with restrained (blue) and unrestrained (green) flip-flop.

At 30 mol% CHOL (Figure 7b), restraining CHOL flip-flop causes pronounced shifts in the DRFs of both bilayers. Specifically, the DRF of the PC bilayer decreases from a strongly positive value (∼0.75) to approximately −0.25, while that of the PE bilayer shifts from a strongly negative value converging toward approximately −0.85 in the unrestrained case to a value near −0.25 when flip-flop is restrained. The snapshots shown in Figures S11 and S12 indicate that, under CHOL flip-flop restraint, lipid domains in both bilayers lose their well-defined transbilayer organization, with registered and anti-registered regions coexisting.

These results demonstrate that lipid headgroup properties and CHOL concentration influence lipid domain registration and anti-registration in a nontrivial and cooperative manner. At 12.5 mol% CHOL, PC bilayer, with their larger and more weakly interacting headgroups, maintains a flat bilayer morphology, such that registered domains persist even when CHOL flip-flop is restrained. In PE bilayers, by contrast, the smaller and more strongly interacting headgroups leave the bilayer closer to the onset of negative spontaneous curvature. Registered domains are still stable under unrestrained conditions, where CHOL flip-flop enhances interleaflet coupling (Figure 2), but restraining flip-flop disrupts this coupling and produces a mixture of registered and anti-registered domains (Figure S10).

At 30 mol% CHOL, CHOL flip-flop becomes strongly suppressed in PE bilayers with smaller and more strongly interacting headgroups, which exhibit anti-registered domains, whereas in PC bilayers with larger and more weakly interacting headgroups, flip-flop is only moderately reduced and registered domains remain stable, as shown in Figures 2 and 5. Once CHOL flip-flop is artificially restrained, however, lipid domains in both bilayers become uncorrelated. These findings indicate that interleaflet coupling mediated by CHOL flip-flop is a key factor governing domain registration and anti-registration at high CHOL concentrations. This applies to both bilayer types, including PE bilayers in which flip-flop is already substantially reduced, where further suppression nonetheless alters the transbilayer organization of the domains.

Previous studies have shown that midplane CHOL, located between the two leaflets of the bilayer, plays an important role in lipid domain registration.^27,28^ Our results are qualitatively consistent with these findings. Most importantly, the present work further reveals that lipid domain registration and anti-registration are cooperatively determined by CHOL concentration and lipid headgroup properties. In our simulations, domains can adopt either registered or anti-registered configurations even when the fraction of midplane CHOL molecules is nearly identical in PC and PE bilayers at a given CHOL concentration (Figure S13), indicating that midplane CHOL abundance alone does not determine the registration state and highlighting the role of headgroup-dependent packing in interleaflet coupling.

The curvature-driven mechanism identified here relates to several established contributions to interleaflet coupling. Conceptually, it parallels diffusional softening, in which the lateral redistribution of curvature-sensitive lipids couples to membrane deformation.^52^ It is also not the only route to anti-registration: Enoki and Heberle observed anti-registered phases in cholesterol-free asymmetric bilayers, attributed to a midplane surface tension penalizing juxtaposed ordered and disordered leaflets,^17^ and Fowler et al. showed that hydrophobic thickness mismatch alone can drive anti-registration, particularly for domains below approximately 20 nm.^13^ Interleaflet coupling therefore reflects multiple contributions, including spontaneous curvature, midplane surface tension, and hydrophobic mismatch, whose relative importance depends on lipid composition, cholesterol content, and domain size. Notably, the PC and PE systems were simulated in identical box dimensions, so finite-size effects on domain organization apply equally to both and cannot account for the headgroup dependence reported here.

## Conclusions

In this work, we used coarse-grained molecular dynamics simulations combined with a FEP-like headgroup-interpolation approach to examine how lipid headgroup properties and CHOL concentration jointly regulate domain registration and anti-registration in symmetric model bilayers. By continuously interpolating between PE-like and PC-like headgroups at a fixed tail composition, we examine the contribution of headgroup properties independently of changes in tail composition and show that their cooperative effects on domain organization are closely linked to changes in membrane curvature and CHOL flip-flop.

In PE-like bilayers, the smaller and more strongly interacting headgroups resist interfacial rearrangement. Consequently, the packing perturbation induced by CHOL incorporation is accommodated primarily through reorganization of the lipid tails, promoting negative membrane curvature and favoring anti-registration. In contrast, PC-like bilayers, with larger and more weakly interacting headgroups, exhibit greater interfacial packing flexibility and less pronounced tail reorganization, allowing the membrane to remain relatively flat and registered over a broader range of CHOL concentrations. CHOL flip-flop provides an additional concentration-dependent contribution to interleaflet coupling. At 12.5 mol% CHOL, rapid flip-flop strengthens this coupling and sustains registration in both bilayer types, whereas its suppression at 30 mol% in PE-like bilayers coincides with the stabilization of anti-registration. Control simulations with restrained CHOL flip-flop result in domain decorrelation in both systems, indicating that flip-flop actively strengthens interleaflet coupling rather than merely accompanying domain registration.

These results identify headgroup-dependent packing, CHOL-induced curvature, and CHOL flip-flop as cooperative determinants of lipid domain organization in symmetric model membranes. The headgroup-interpolation strategy further provides a controlled framework for dissecting lipid-specific contributions to interleaflet coupling in more complex membrane systems.

## Author Contributions

J. C. and Y. C.: Conceptualization, Investigation, Formal Analysis, and Writing – Original Draft. These authors contributed equally to this work. D. P. T.: Supervision and Writing – Review & Editing. Q. L.: Conceptualization, Supervision, and Writing – Review & Editing.

## Notes

The authors declare no competing financial interest.

## Supporting Information

The Supporting Information is available free of charge.

- Representative snapshots of phase separation processes are provided for both CHOL-unrestrained lipid bilayers and CHOL-restrained lipid bilayers, Lo-domain surface areas, CHOL distributions, and leaflet imbalance, lipid-tail order parameters calculated relative to local membrane normals, and the fraction of midplane CHOL.

## Acknowledgement

1. J. C. and Q. L. thank Dr. Yong Wang (Zhejiang University) for helpful discussions. This work is supported by the Zhejiang Provincial Natural Science Foundation of China under Grant No. LZ25A040005, and by the Natural Sciences and Engineering Research Council of Canada (DPT). This research was undertaken, in part, thanks to funding from the Canada Research Chairs program. Computational resources provided by Compute Canada are appreciated.

## Notes

### Competing Interest Statement

The authors have declared no competing interest.

### Summary of Updates

This version has been substantially revised to improve the analysis, presentation, and mechanistic interpretation of the results. We expanded the discussion of how cholesterol concentration and lipid headgroup chemistry regulate interleaflet domain registration, and further clarified the roles of lipid packing, membrane curvature, cholesterol flip-flop, and interleaflet coupling. Several figures and corresponding sections of the manuscript have also been revised and reorganized for clarity.

