## Supporting Information for "Cooperative Regulation of Membrane Domain Registration by Lipid Headgroup Properties and Cholesterol Dynamics"

### These authors contributed equally.

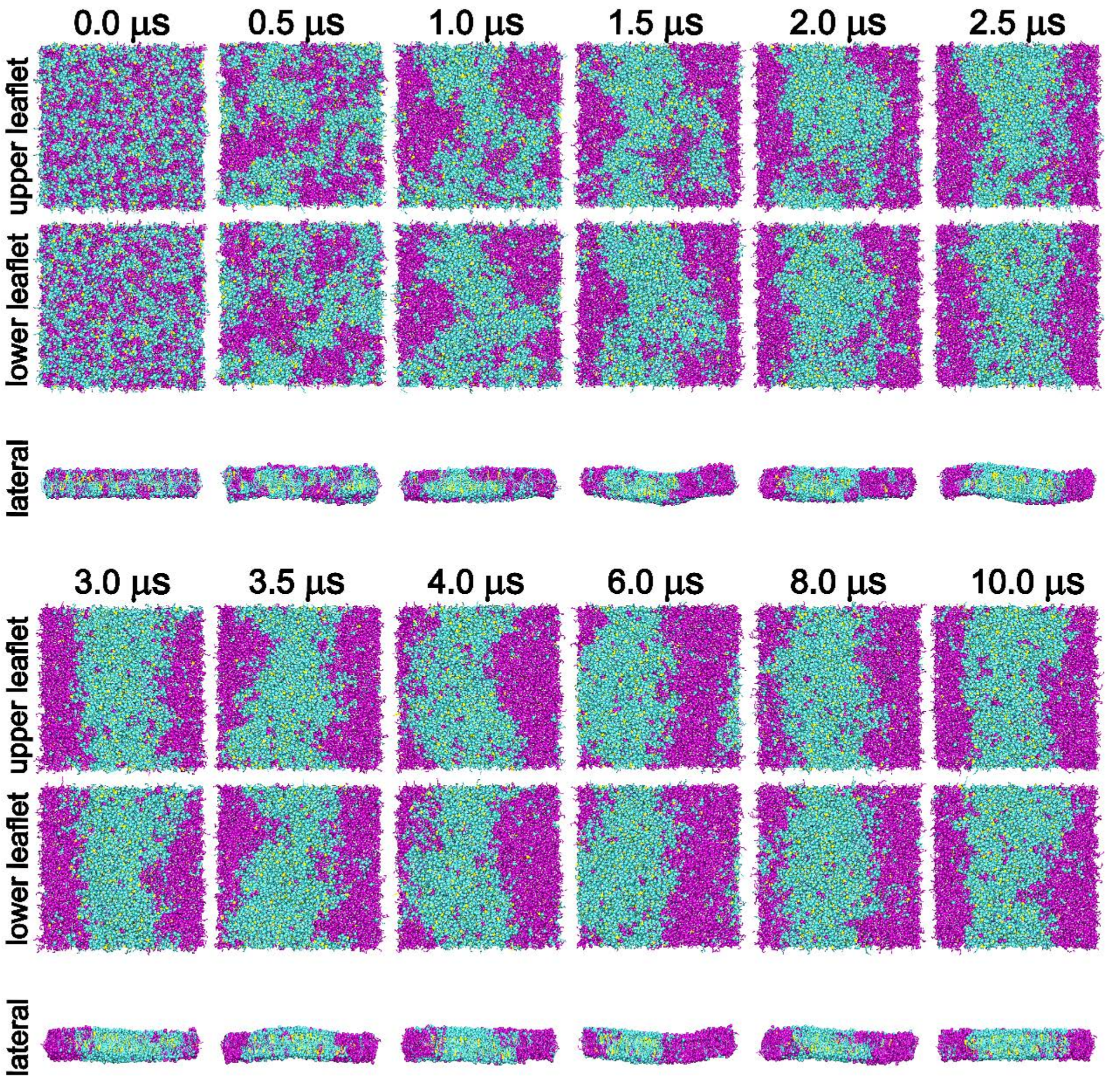


**Figure S1.** Representative snapshots of the phase separation process in a lipid bilayer composed of DPPC, DIPC, and CHOL at a molar ratio of 4:3:1 (12.5 mol% CHOL). DPPC, DIPC, and CHOL molecules are shown in cyan, magenta, and yellow, respectively.

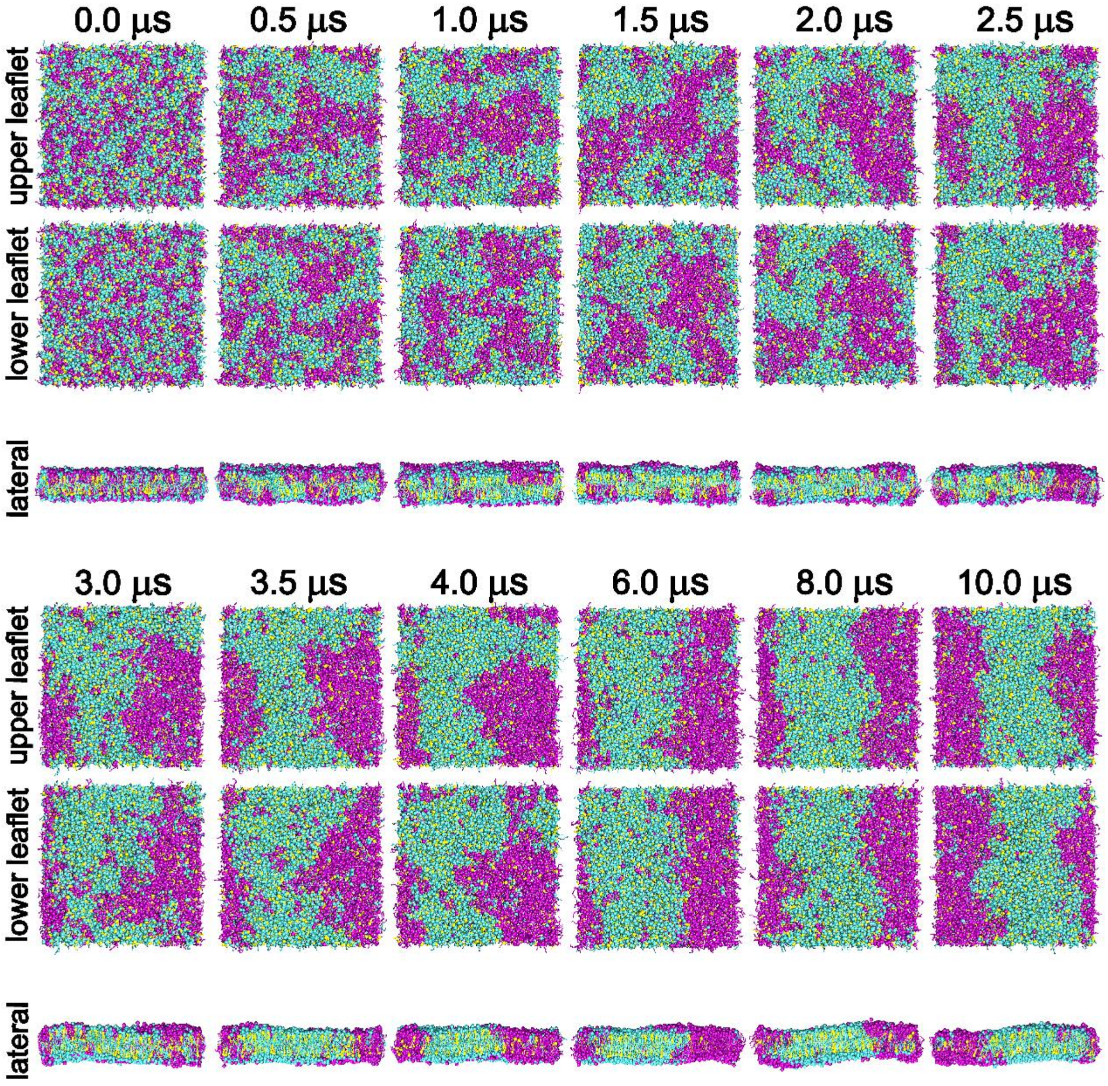


**Figure S2.** Representative snapshots of the phase separation process in a lipid bilayer composed of DPPC, DIPC, and CHOL at a molar ratio of 4:3:3 (30 mol% CHOL). DPPC, DIPC, and CHOL molecules are shown in cyan, magenta, and yellow, respectively.

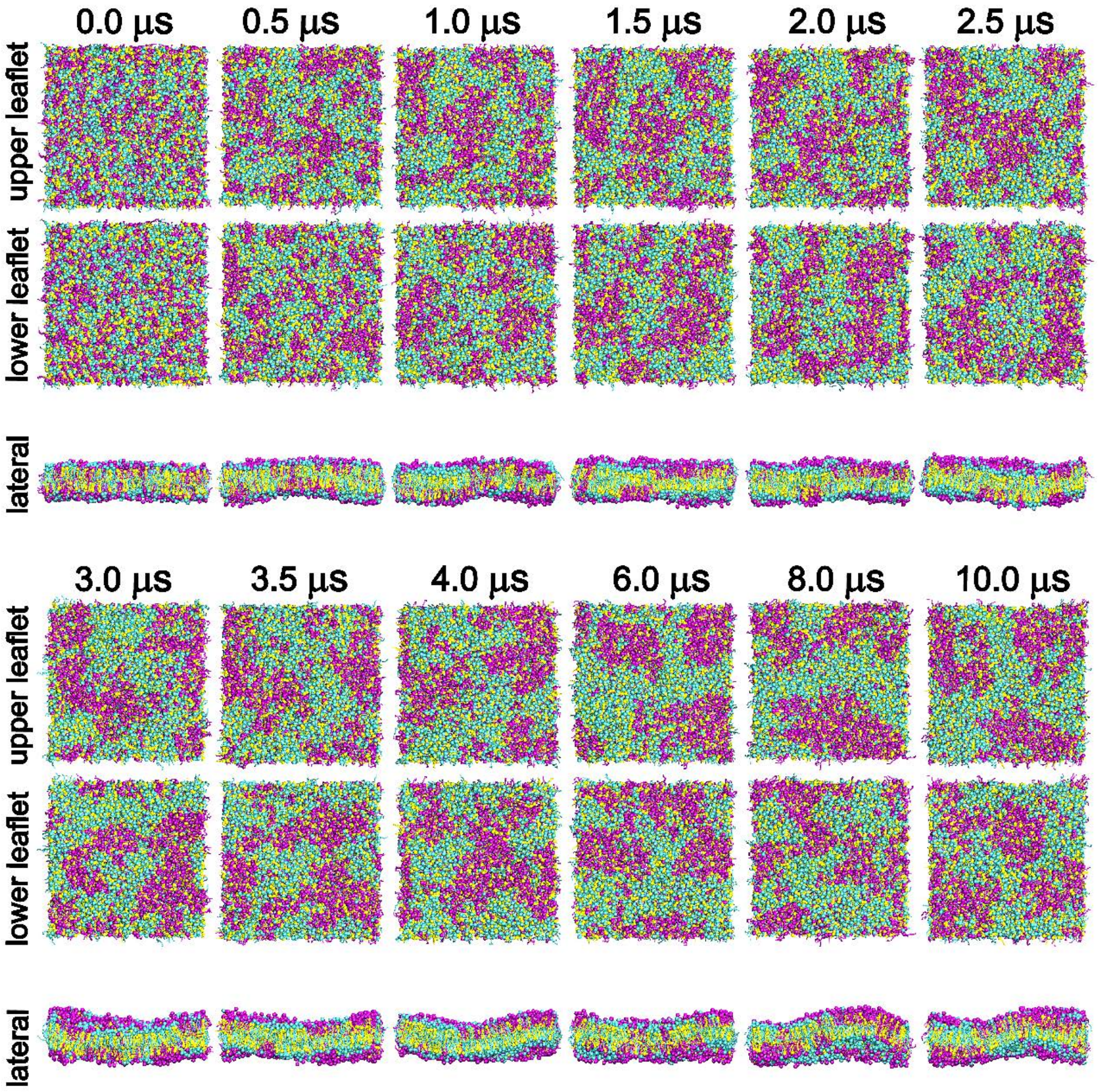


**Figure S3.** Representative snapshots of the phase separation process in a lipid bilayer composed of DPPC, DIPC, and CHOL at a molar ratio of 4:3:7 (50 mol% CHOL). DPPC, DIPC, and CHOL molecules are shown in cyan, magenta, and yellow, respectively.

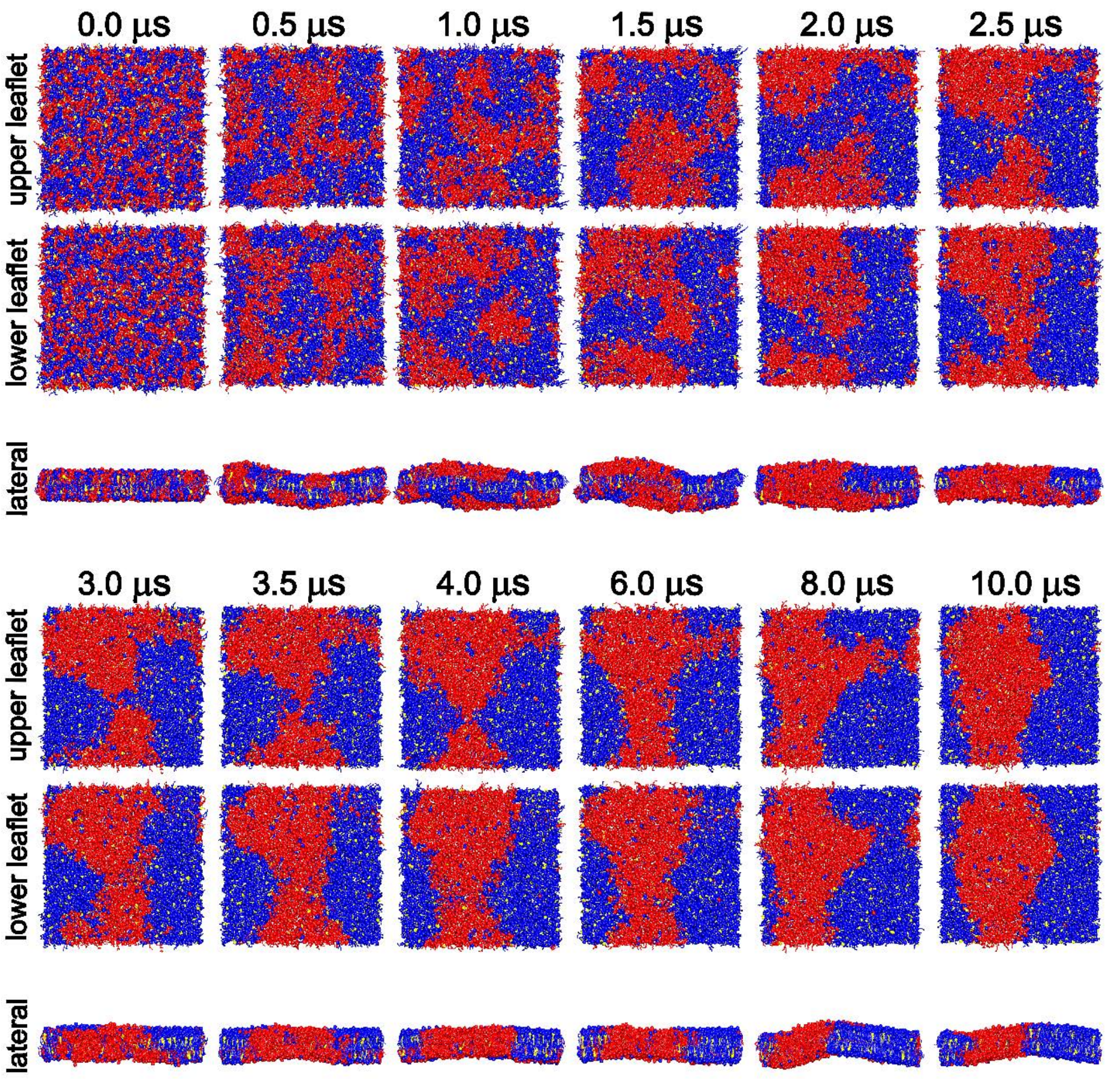


**Figure S4.** Representative snapshots of the phase separation process in a lipid bilayer composed of DPPE, DIPE, and CHOL at a molar ratio of 4:3:1 (12.5 mol% CHOL). DPPE, DIPE, and CHOL molecules are shown in blue, red, and yellow, respectively.

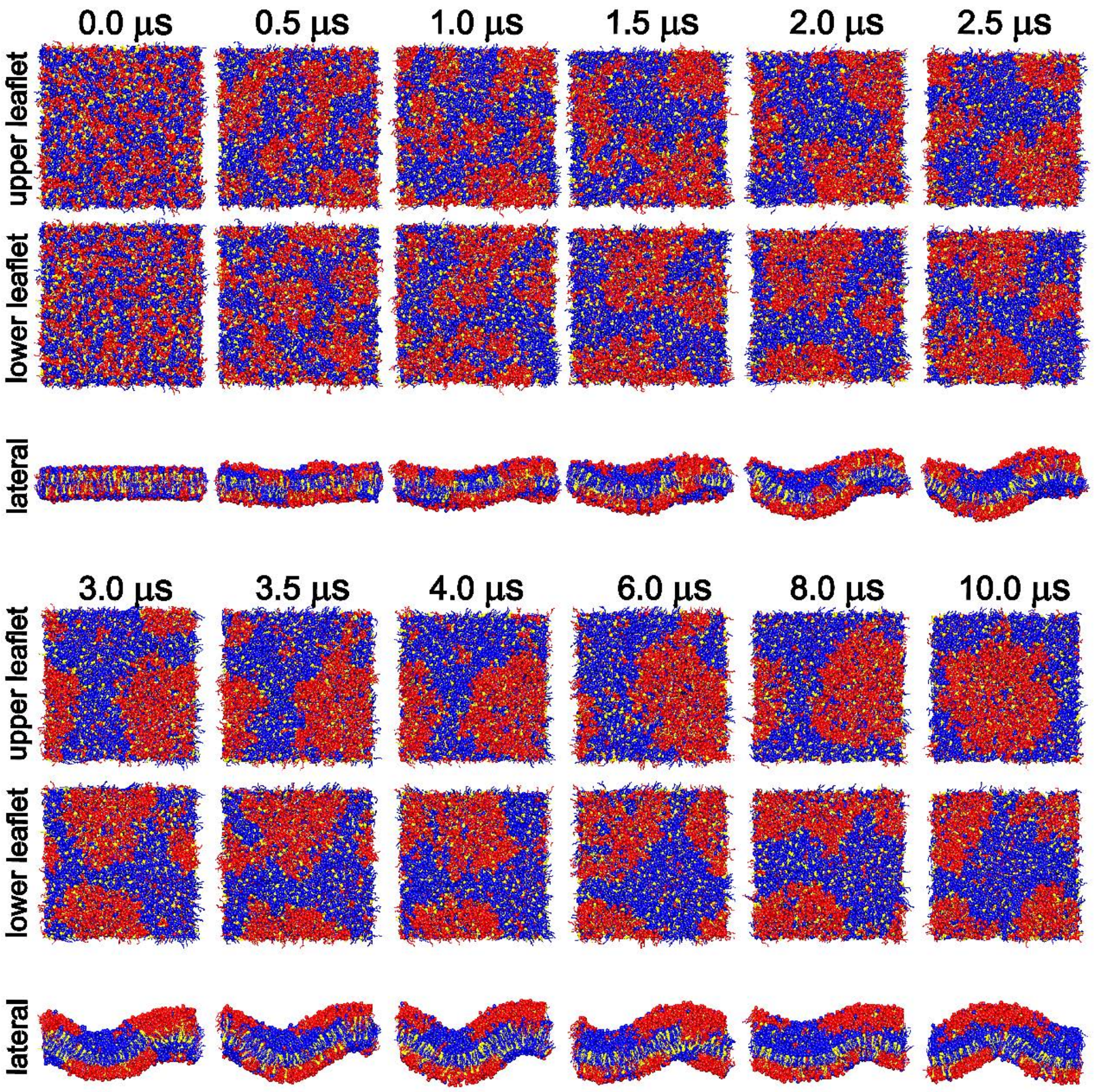


**Figure S5.** Representative snapshots of the phase separation process in a lipid bilayer composed of DPPE, DIPE, and CHOL at a molar ratio of 4:3:3 (30 mol% CHOL). DPPE, DIPE, and CHOL molecules are shown in blue, red, and yellow, respectively.

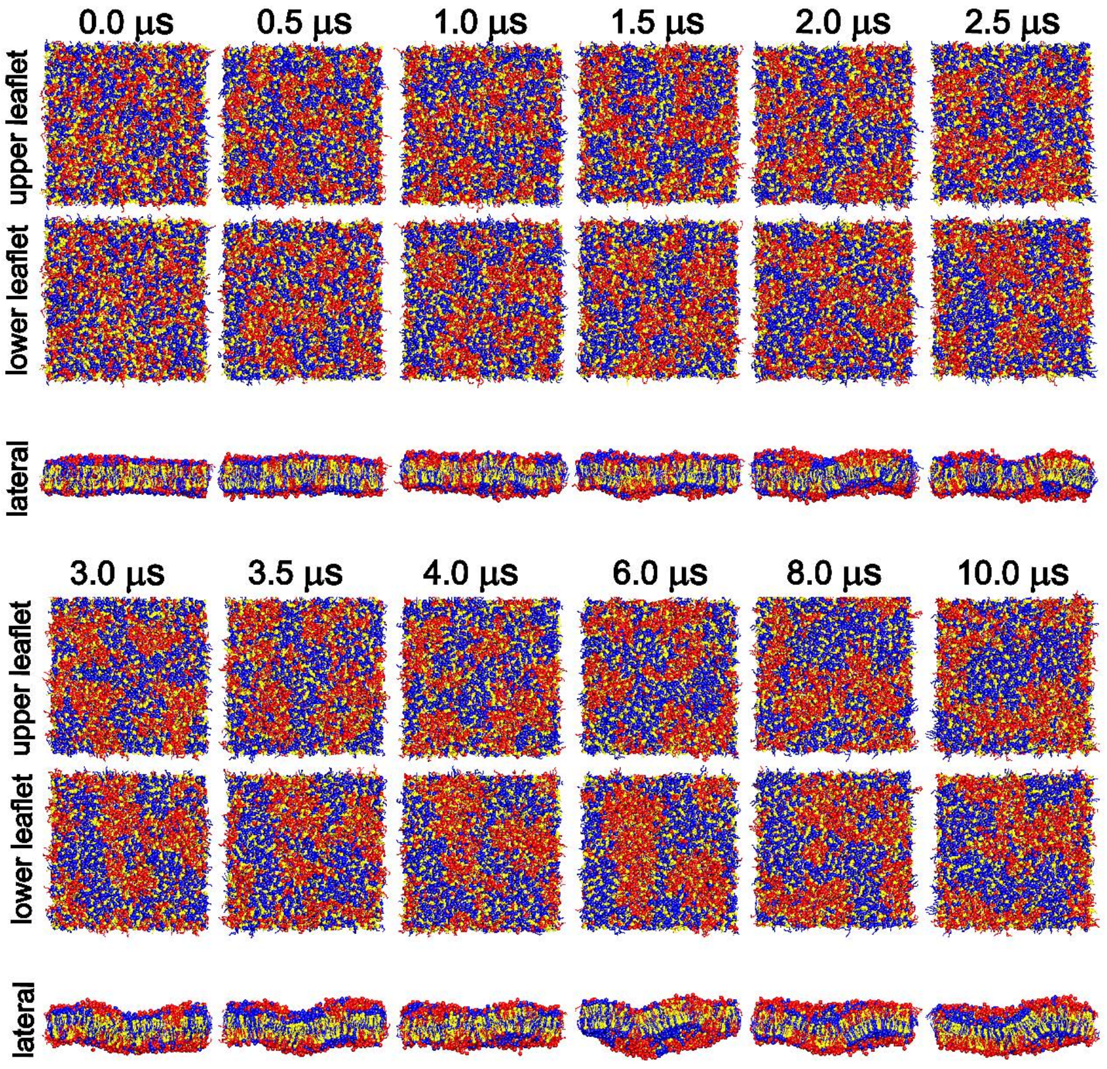


**Figure S6.** Representative snapshots of the phase separation process in a lipid bilayer composed of DPPE, DIPE, and CHOL at a molar ratio of 4:3:7 (50 mol% CHOL). DPPE, DIPE, and CHOL molecules are shown in blue, red, and yellow, respectively.

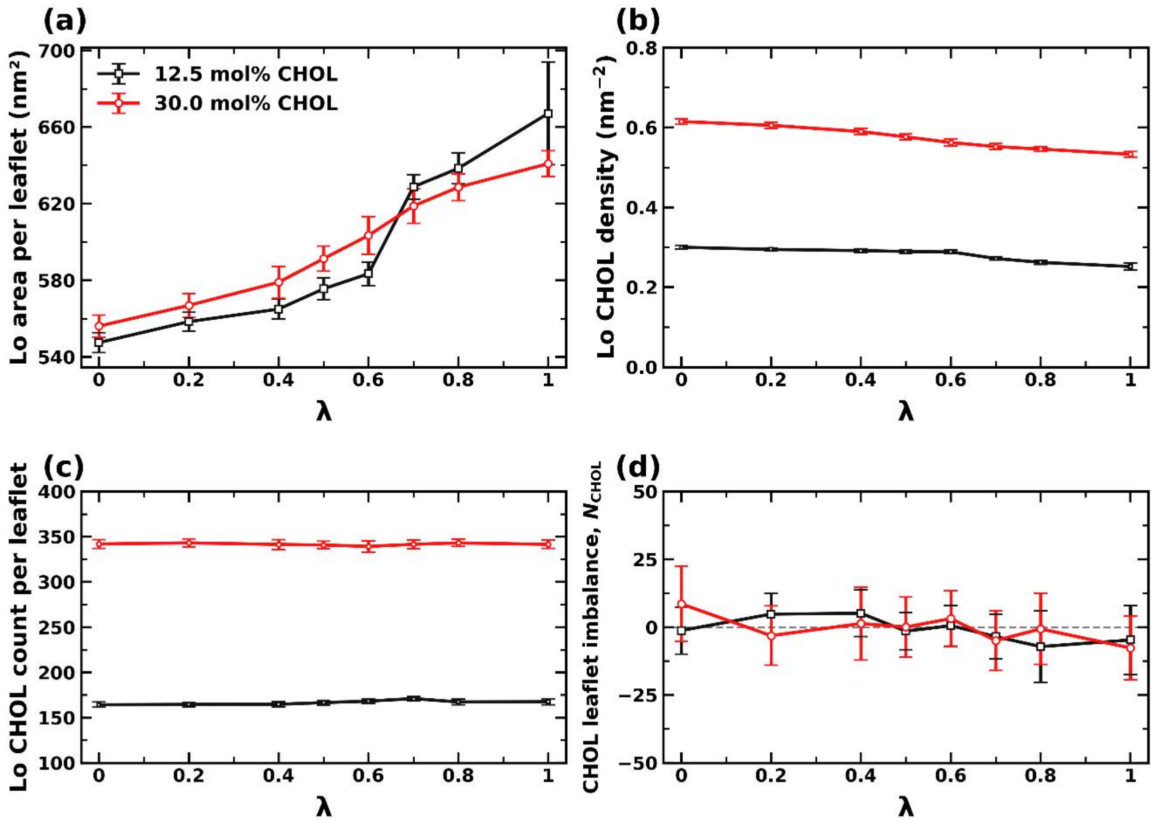


**Figure S7.** (a) Mean Lo-domain surface area per leaflet, (b) mean CHOL number density within the Lo domains, and (c) mean number of CHOL molecules within the Lo domains as functions of λ for bilayers containing 12.5 mol% CHOL (black squares) and 30.0 mol% CHOL (red circles). For panels (a–c), the upper- and lower-leaflet values were averaged, and all Lo domains present in each leaflet were included. (d) Difference in the total CHOL counts between the upper and lower leaflets, defined as $\Delta N_{\mathrm{CHOL}} = N_{\mathrm{CHOL,upper}} - N_{\mathrm{CHOL,lower}}$. All values were averaged over 8–10 μs, and error bars indicate the standard deviations over this time interval.

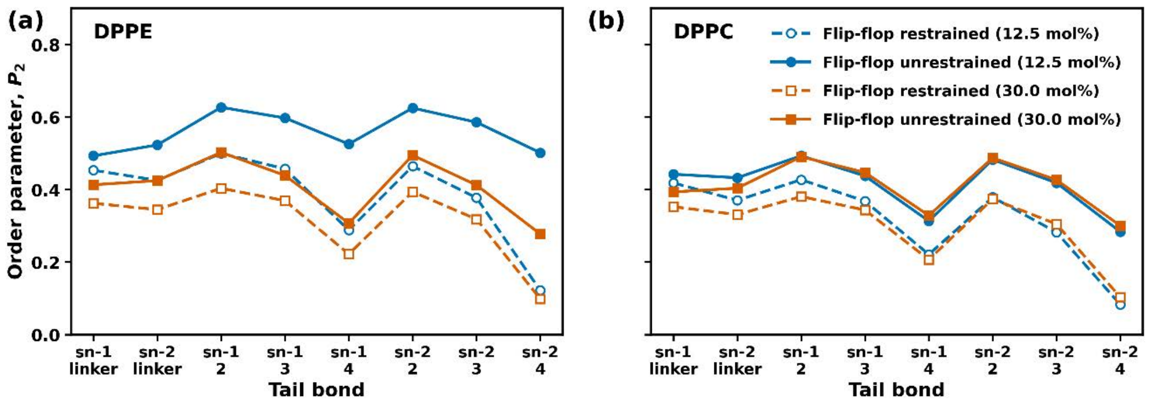


**Figure S8.** Lipid-tail order parameters ($P_2$) calculated relative to local membrane normals over the final 5 μs for (a) DPPE and (b) DPPC. Results are shown for bilayers with 12.5 mol% CHOL (blue) and 30 mol% CHOL (orange), comparing simulations with restrained CHOL flip-flop (open symbols and dashed lines) and the corresponding simulations with unrestrained CHOL flip-flop (filled symbols and solid lines). Values are presented for the linker and successive bonds along the sn-1 and sn-2 lipid tails.

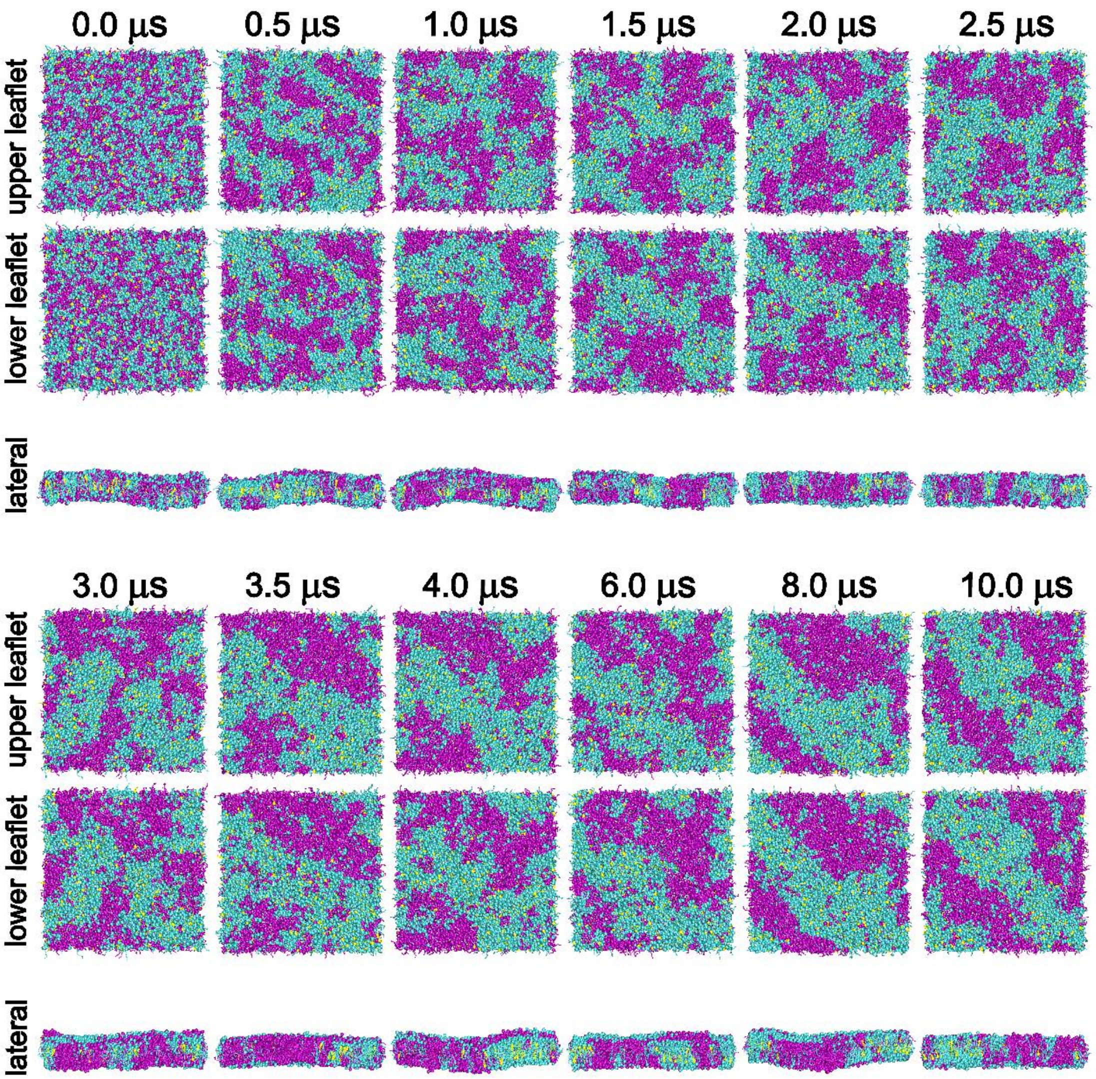


**Figure S9.** Representative snapshots of the phase separation process in a lipid bilayer composed of DPPC, DIPC, and CHOL at a molar ratio of 4:3:1 (12.5 mol% CHOL) with restrained CHOL flip-flop. DPPC, DIPC, and CHOL molecules are shown in cyan, magenta, and yellow, respectively.

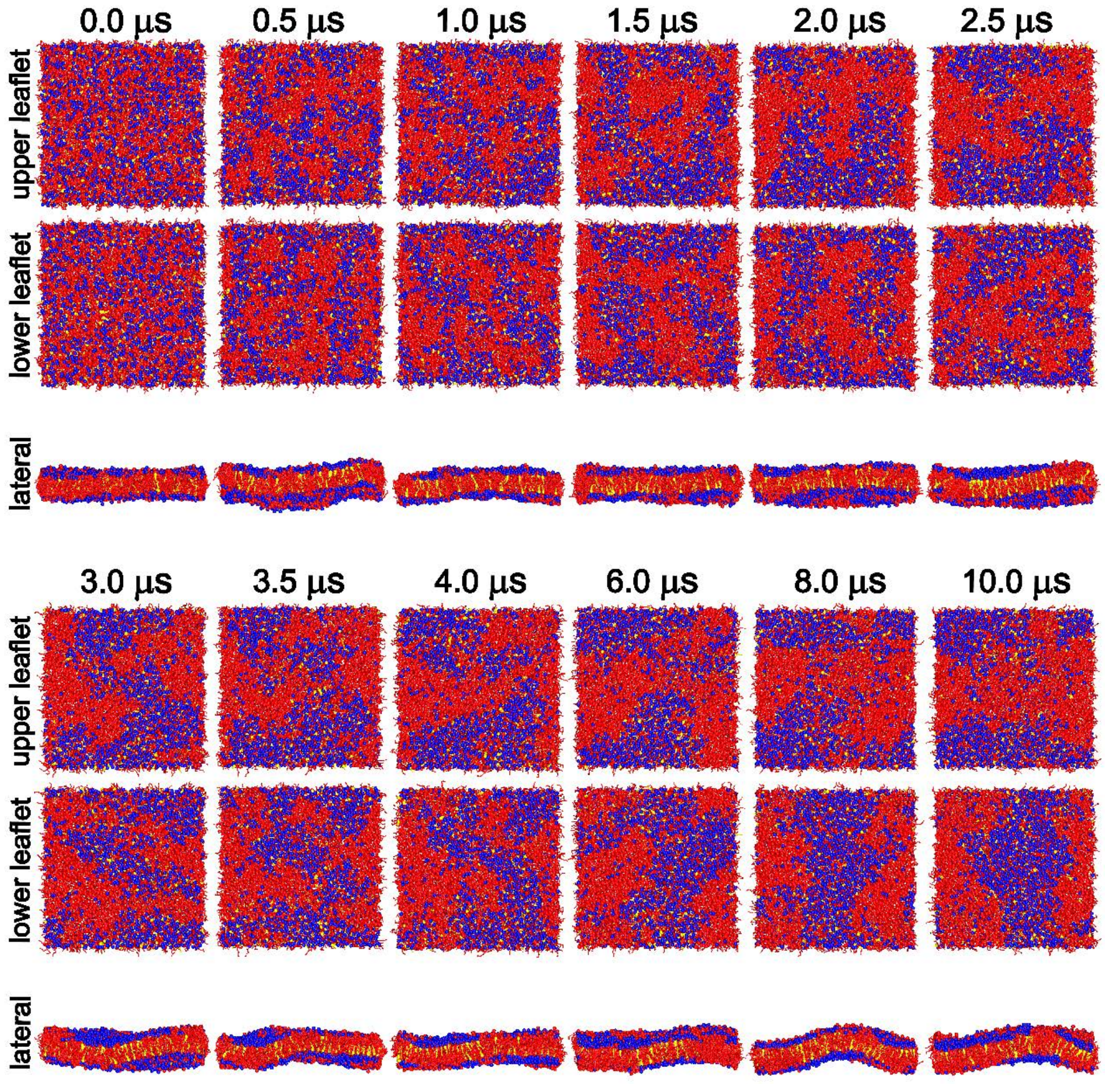


**Figure S10.** Representative snapshots of the phase separation process in a lipid bilayer composed of DPPE, DIPE, and CHOL at a molar ratio of 4:3:1 (12.5 mol% CHOL) with restrained CHOL flip-flop. DPPE, DIPE, and CHOL molecules are shown in blue, red, and yellow, respectively.

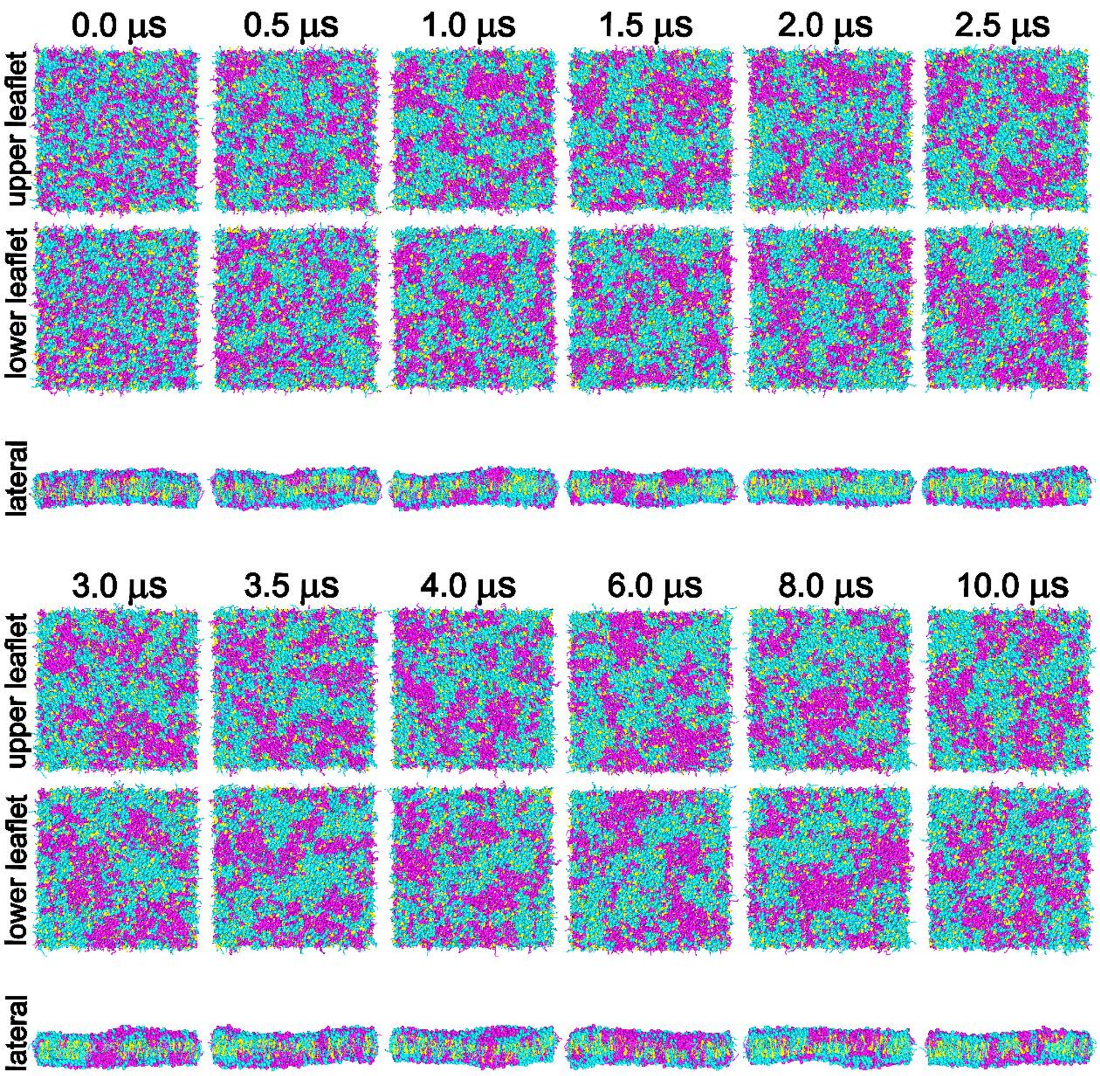


**Figure S11.** Representative snapshots of the phase separation process in a lipid bilayer composed of DPPC, DIPC, and CHOL at a molar ratio of 4:3:3 (30 mol% CHOL) with restrained CHOL flip-flop. DPPC, DIPC, and CHOL molecules are shown in cyan, magenta, and yellow, respectively.

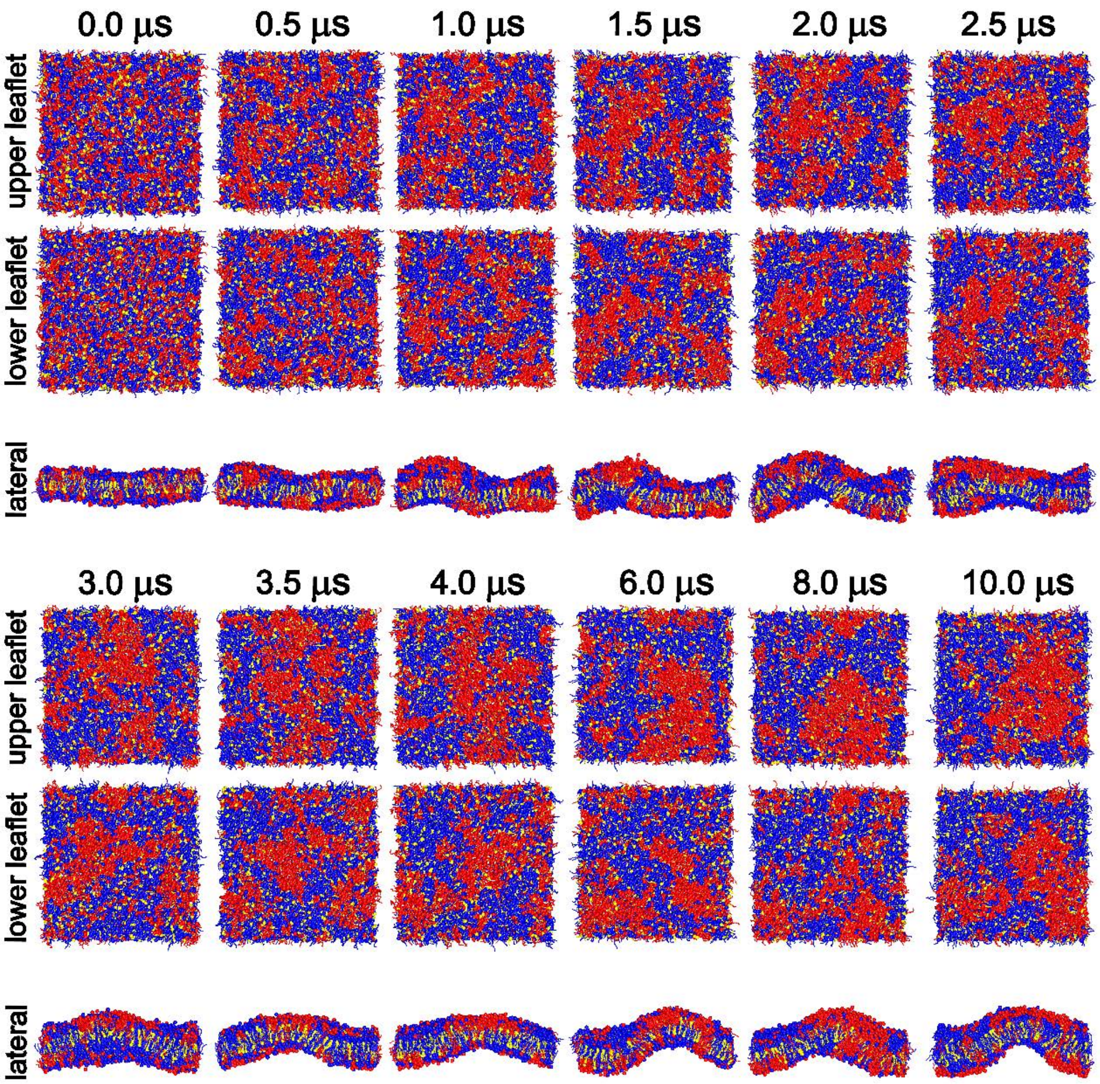


**Figure S12.** Representative snapshots of the phase separation process in a lipid bilayer composed of DPPE, DIPE, and CHOL at a molar ratio of 4:3:3 (30 mol% CHOL) with restrained CHOL flip-flop. DPPE, DIPE, and CHOL molecules are shown in blue, red, and yellow, respectively.

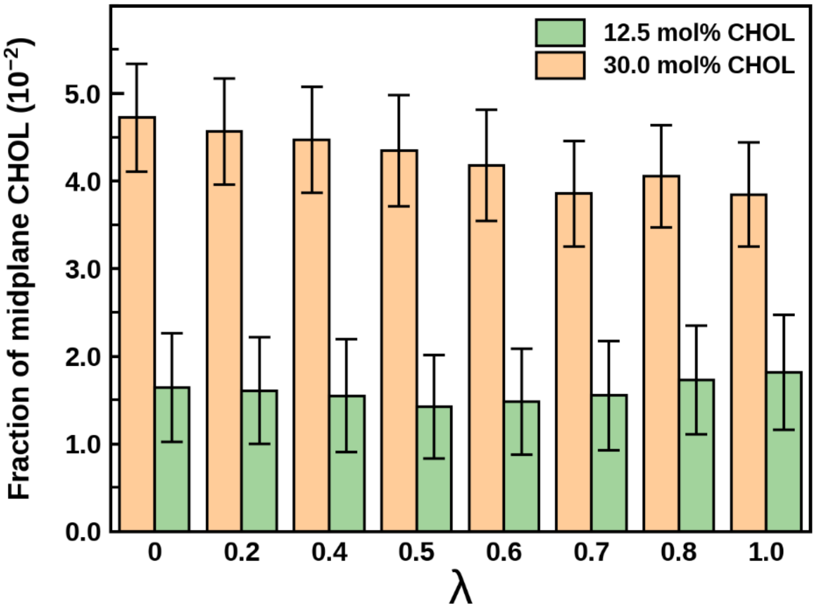


**Figure S13.** Time-averaged fraction of midplane CHOL molecules as a function of λ in the eight FEP-like simulation replicas at 12.5 and 30 mol% CHOL. The coupling parameter λ varies from 0 (PE-like headgroups) to 1 (PC-like headgroups). Error bars indicate the standard deviation over the analyzed trajectory segment from 5 to 10 μs.
